# Mesenchymal tumor organoid models recapitulate rhabdomyosarcoma subtypes

**DOI:** 10.1101/2022.01.03.474504

**Authors:** Michael T. Meister, Marian J. A. Groot Koerkamp, Terezinha de Souza, Willemijn B. Breunis, Ewa Frazer-Mendelewska, Mariël Brok, Jeff DeMartino, Freek Manders, Camilla Calandrini, Hinri H. D. Kerstens, Alex Janse, M. Emmy M. Dolman, Selma Eising, Karin P. S. Langenberg, Marc van Tuil, Rutger R. G. Knops, Sheila Terwisscha van Scheltinga, Laura S. Hiemcke-Jiwa, Uta Flucke, Johannes H. M. Merks, Max M. van Noesel, Bastiaan B. J. Tops, Jayne Y. Hehir-Kwa, Patrick Kemmeren, Jan J. Molenaar, Marc van de Wetering, Ruben van Boxtel, Jarno Drost, Frank C. P. Holstege

## Abstract

Rhabdomyosarcomas (RMS) are mesenchyme-derived tumors and the most common childhood soft tissue sarcomas. Treatment is intense, with a nevertheless poor prognosis for high-risk patients. Discovery of new therapies would benefit from additional preclinical models. Here we describe the generation of a collection of pediatric RMS tumor organoid (tumoroid) models comprising all major subtypes. For aggressive tumors, tumoroid models can often be established within four to eight weeks, indicating the feasibility of personalized drug screening. Molecular, genetic and histological characterization show that the models closely resemble the original tumors, with genetic stability over extended culture periods of up to six months. Importantly, drug screening reflects established sensitivities and the models can be modified by CRISPR/Cas9 with *TP53* knockout in an embryonal RMS model resulting in replicative stress drug sensitivity. Tumors of mesenchymal origin can therefore be used to generate organoid models, relevant for a variety of preclinical and clinical research questions.

## Introduction

Rhabdomyosarcoma (RMS) is a type of malignant tumor of mesenchymal origin (Yang et al., 2014) and forms the most common soft tissue sarcoma in children and adolescents (Li et al., 2008). Historically, RMS has been divided into two main subtypes based on histology. Whereas embryonal RMS (eRMS) displays cellular heterogeneity and hallmarks of immature skeletal myoblasts (Patton and Horn, 1962), alveolar RMS (aRMS) cells are distributed around an open central space, thereby resembling pulmonary alveoli (Enterline and Horn, 1958). eRMS is more frequently observed in children under ten, accounting for two-thirds of all RMS cases, and generally has a better prognosis than aRMS, which is more common in adolescents and young adults (Perez et al., 2011). In aRMS, a sole genetic driver alteration is usually observed, caused by a chromosomal translocation resulting in a fusion gene between either *PAX3* or *PAX7* and *FOXO1*. In contrast, eRMS is genetically more heterogeneous, harboring mutations in several common oncogenes or tumor suppressor genes (Shern et al., 2014). Other subtypes of RMS have recently been recognized (WHO, 2020). RMS treatment is guided by protocols developed by multi-national collaborative groups and includes systemic chemotherapy in addition to local therapy (radiotherapy and/or surgery) (Skapek et al., 2019). The prognosis of RMS has improved over the last decades (Bisogno et al., 2019). For patients with high-risk, refractory or relapsed disease, prognosis remains poor however, despite an immense treatment burden (Mascarenhas et al., 2019; Pappo et al., 1999). Thus, development of new therapeutic options is of critical importance for these patients.

Development of such treatment options requires *in vitro* models and may therefore benefit from application of organoid technology. The basis of this technology is that given a suitable growth environment, tissue stem cells self-renew as well as give rise to natural progeny which organize according to their preferred growth modality without the need for artificial cell immortalization. The technology was first established in healthy epithelial tissue from mouse small intestine (Sato et al., 2009) and soon adapted to various other healthy and diseased epithelial tissues, including cancer (Clevers, 2016). Tumor organoid (tumoroid) systems are proving useful in cancer research as they display genetic stability over extended culture periods, retaining the molecular characteristics of the tumor they are derived from. Tumoroid models can be expanded, facilitating high-throughput screening approaches such as small molecule or CRISPR/Cas9-knockout screening (Bleijs et al., 2019).

To date, tumoroid approaches have been primarily applied to cancers derived from epithelial cells (i.e., carcinomas). Recent studies demonstrate that deriving tumoroid models from non-epithelial cancer is feasible but this has as yet not been achieved for pure mesenchymal cancers (Abdullah et al., 2021; Fusco et al., 2019; Jacob et al., 2020; Saltsman et al., 2020; Yamazaki et al., 2021). Application to tumors of mesenchymal origin such as RMS would be of obvious benefit. Tumoroid models of pediatric nephroblastoma (Wilms tumors) have been described, which, depending on the subtype, can contain stromal cells (Calandrini et al., 2020). In addition, cells derived from synovial sarcoma and other adult soft tissue sarcomas can grow to a limited extent on fetal calf serum, which, although undefined in terms of the required essential growth factors, also indicates feasibility (Boulay et al., 2021; Brodin et al., 2019). Furthermore, *in vitro* propagation of RMS tumor cells derived from patient-derived xenograft (PDX) mouse models has recently been shown (Manzella et al., 2020). Although these results are encouraging, no directly patient-derived collection of tumoroid models of malignant tumors of pure mesenchymal origin (i.e., sarcomas) has been generated and studied after growth for extensive periods in well-defined media components. In this study, we therefore set out to develop and apply approaches for generating a collection of tumoroid models that covers the major RMS subtypes, a pediatric cancer of mesenchymal origin with poor outcome for high-risk patients. Besides generating and extensively characterizing the tumoroid collection, we also investigated applicability for drug screening and genetic modification (Fig. 1A).

**Fig. 1.**
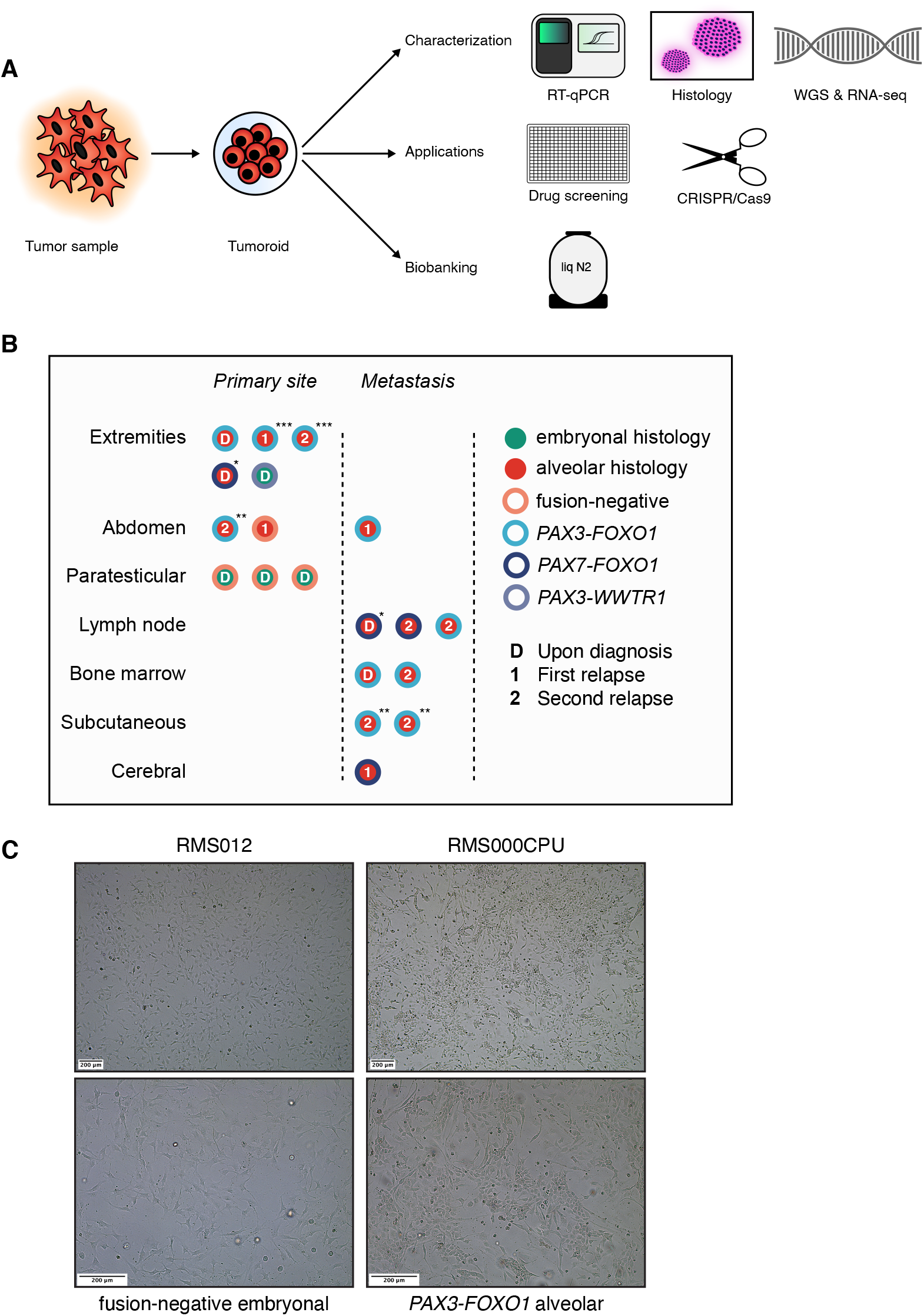
A collection of RMS tumoroid models that represent the diverse clinical presentation of RMS: (A) Tumor organoid (tumoroid) pipeline. WGS = whole-genome sequencing, RNA-seq = mRNA sequencing, liq N2 = liquid nitrogen. (B) Overview of available RMS tumoroid models in the collection separated by primary vs. metastatic site and exact tumor location. The color of the inner circle indicates the histological subtype while the color of the outer circle indicates the presence or absence of a fusion transcript. Letters within the circle indicate disease instance. Asterisks mark tumoroid models derived from the same patient but from distinct tumor samples. (C) Brightfield microscopy images of two representative RMS tumoroid models from a fusion-negative embryonal and a *PAX3-FOXO1* fusion-positive alveolar tumoroid model grown in a two-dimensional monolayer in two magnifications as indicated by the scale bars.

## Results

### A protocol to collect and process RMS tumor samples for tumoroid model establishment and propagation

Before starting to generate a collection of RMS tumoroid models, we first optimized sample acquisition and logistics between surgery, pathology and organoid culture labs (Materials & Methods). In parallel to optimizing sample acquisition, we also optimized sample processing, including testing different formulations of growth media by a combination of systematic and trial and error approaches (Discussion). RMS tumor samples are diverse. Most samples are small needle biopsies (i.e, 16-gauge tru-cut), as large resection specimens are mostly restricted to pre-treated RMS or to treatment-naïve paratesticular fusion-negative eRMS (FN-eRMS). In addition, a subset of samples (4% here) are not solid, being acquired as bone marrow aspirates of infiltrating tumor cells (Fig. 1B). Samples are plated as minced pieces embedded in a droplet of extracellular matrix (ECM) substitute (Basement-Membrane Extract, BME) and as single-cell suspensions in BME-supplemented medium. Outgrowth of tumor cells to tumoroid models can occur from both modalities. In the case of successful outgrowth of initially plated cells, cells organize as two-dimensional monolayers (Fig. 1C). This appears to be the cells’ preferred growth modality, as plating them as single-cell suspensions in BME droplets results in cells escaping the surrounding matrix and sinking to the bottom of the culture plate from which they continue to grow in a monolayer. Therefore, cells are further propagated and expanded in this way. We considered an RMS tumoroid model to be successfully established if, over the course of culturing, the expression of specific tumor markers is retained and the culture expansion is at least sufficient for drug screening, all as described below.

### Early detection of tumor cells during culturing

Tumors consist of a variety of different cell types. These include normal cell types that can grow as well or even better in the provided culture conditions, possibly outcompeting tumor cells (Dijkstra et al., 2020). It would therefore be useful to test for the presence of tumor cells early during culturing to omit the unnecessary propagation of cultures lacking any. At early time points, material is limited, impacting the range of applicable assays. The establishment protocol therefore utilizes an RT-qPCR assay after the first or second passage of cells with probes for standard RMS histopathology markers, i.e., *DES, MYOG, MYOD1* (WHO, 2020), and the fusion transcript in fusion-positive RMS (FP-RMS) (Ponce-Castañeda et al., 2014). We considered a sample positive for tumor cells if at least one of the three genes, plus for FP-RMS the fusion transcript, test positive. All samples that successfully yield tumoroid models, show positivity for at least one marker gene at this stage, while most models (17 out of 19) are positive for all three marker genes and the fusion transcript if applicable (Fig. 2A and B). The RT-qPCR-based approach is therefore a useful tool to determine feasibility at an early stage.

**Fig. 2.**
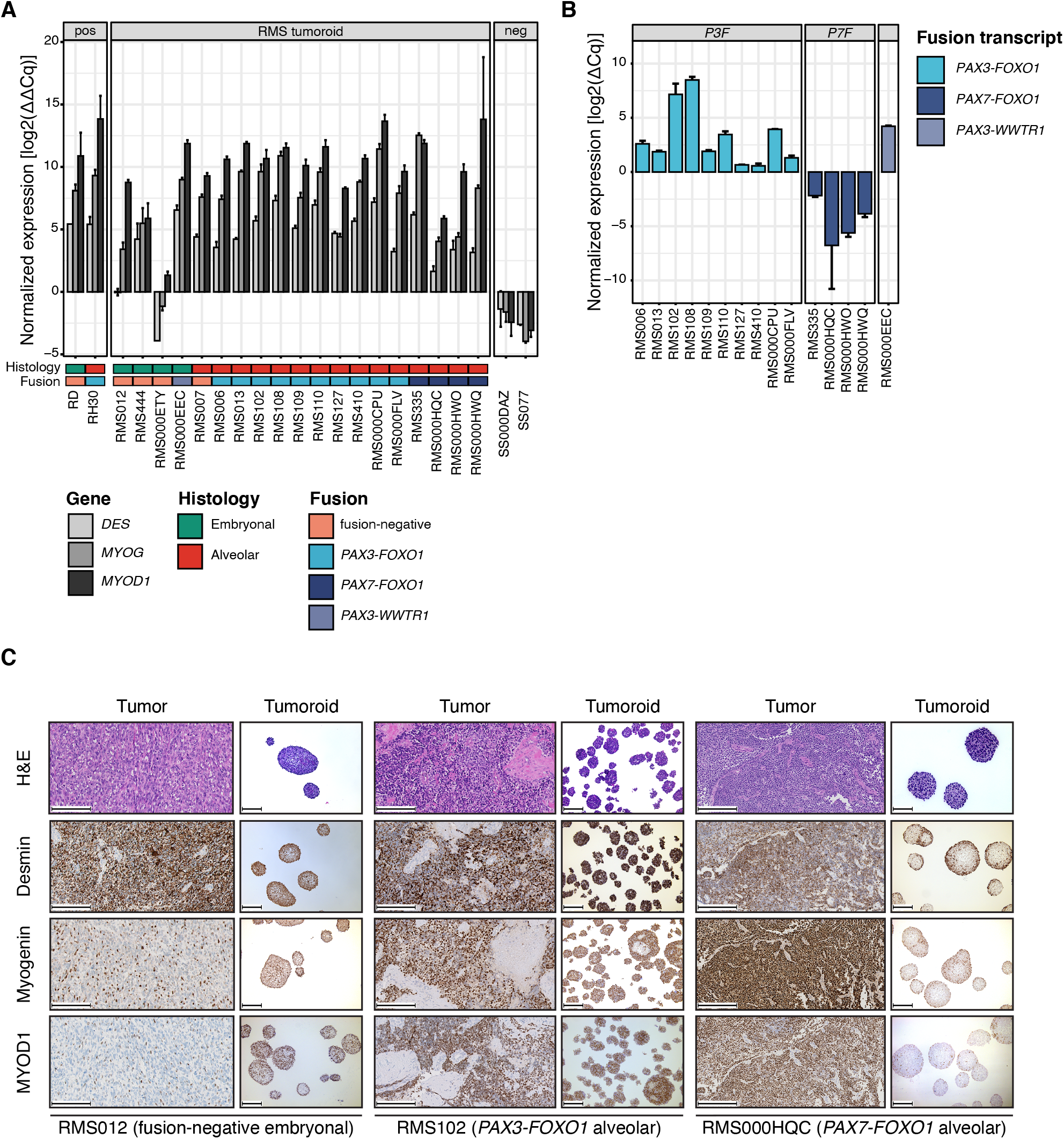
Early detection of tumor cells during culturing and retained marker protein expression: (A) RT-qPCR of early passage RMS tumoroid models shows positivity for at least one gene used in standard-of-care pathology analysis (*DES*, *MYOG*, or *MYOD1*). Conventional RMS cell lines (RD and RH30) were used as positive controls, while two Synovial Sarcoma (SS000DAZ and SS077) tumoroid models were used as negative controls. Gene expression was normalized to the expression of a house-keeping gene and human reference RNA (HREF) via the ΔΔCq method. (B) RT-qPCR of early passage RMS tumoroid models reliably detects the aberrant fusion transcripts. Fusion gene expression was normalized to the expression of a house-keeping gene via the ΔCq method. (C) Morphological (via H&E) and immunohistochemical (IHC) comparison of RMS tumors and derived RMS tumoroid models shows retained marker protein (Desmin, Myogenin and MYOD1) expression and cellular morphology. Scale bars equal 200 µm.

### RMS tumoroid models retain marker protein expression

A hallmark of RMS tumors is the expression of proteins associated with non-terminally differentiated muscle (i.e., Desmin, Myogenin and MYOD1). Expression of these proteins differs between RMS subtypes (Dias et al., 2000) and can be associated with prognosis (Heerema-Mckenney et al., 2008). To properly reflect the original tumors, RMS tumoroid models should therefore retain the expression patterns of these proteins. The RMS tumoroid establishment protocol therefore includes a morphological (H&E) and immunohistochemical (IHC) assessment at the time of successful establishment (i.e., drug screening). To enable comparison between tumoroid models and the tumors they were derived from, models are grown as three-dimensional spheres to mimic the three-dimensional architecture of tumors (Fig. 2C). When evaluated in this manner, RMS tumoroid models show retained expression levels and patterns of Desmin, Myogenin and MYOD1 at the time of successful establishment. Cellular morphology is also retained, with aRMS models displaying homogenous, primitive cells, with large nuclei and minimal cytoplasm, and eRMS models displaying more heterogenous tumor cells with variable maturation (Fig. 2C and Suppl. Fig. 1), as in the tumors (WHO, 2020). In summary, RMS tumoroid models retain histopathological hallmarks of RMS tumors, giving a first indication that they reflect the tumors they were derived from to a high extent.

### A collection of RMS tumoroid models that represent the diverse clinical presentation of RMS

Having established protocols for the acquisition, processing, initial growth, and characterization of RMS tumoroid models, we applied this to 46 consecutive samples from pediatric RMS patients treated in the Netherlands, resulting in a collection of 19 RMS tumoroid models (41% efficiency). These models are derived from tumors comprising both main histological subtypes (embryonal and alveolar), different fusion types (fusion positive *PAX3-FOXO1, PAX7-FOXO1*, a novel fusion *PAX3-WWTR1*, as well as fusion negative), various risk groups, locations, and are derived from primary as well as relapsed disease (Fig. 1B and Suppl. Table 1). Outgrowth of cells from highly aggressive RMS subtypes shows a higher success rate, as indicated by 83% successful establishment for FP-RMS versus 16% for FN-RMS, and 61% success in relapsed, versus 30% in primary RMS tumors, respectively. There was no statistically significant difference in the success rate between recently treated and untreated tumors (33% versus 43%, respectively, *p* = 0.59, chi-square test). Similar differences between relapsed and primary tumors in establishment rates have been observed for orthotopic PDX models (O-PDX) of RMS (Stewart et al., 2017). Compared to O-PDX models of RMS, the overall success rate is lower for establishing RMS tumoroid models (41% versus 65%, respectively). While the success rate is lower, RMS tumoroid models can be more rapidly established. Engraftment of O-PDX RMS models takes 1 to 5.5 months before tumor growth is first observed in mice, while RMS tumoroid models can be fully established, i.e., characterized and subjected to drug screening, in as little as 27 days for highly aggressive tumors. Overall, the median time from acquisition of the tumor sample to successful drug screening was 81 days (with 7 models being screened in less than 2 months). This indicates that our approach could potentially be applied in a personalized medicine setting where it is crucial to obtain results as fast as possible to provide information on treatment options.

A subset of the tumoroid models were derived from the same patient, but at different points during treatment and/or from different body sites (marked with one or more asterisks in Fig. 1B). This potentially facilitates studies of tumor evolution or acquired treatment resistance. Furthermore, the collection contains a model of an eRMS with a novel fusion between *PAX3* and *WWTR1*. Such fusions have been reported as rare events in biphenotypic sinonasal sarcomas, which usually harbor *PAX3-MAML3* fusions (Loarer et al., 2019). Taken together, the protocol efficiently yields tumoroid models from highly aggressive as well as from extremely rare RMS subtypes, resulting in an initial collection covering a broad spectrum of subtypes.

### RMS tumoroid models molecularly resemble the tumor they are derived from

Given that the present tumoroid collection is the first to be established from tumors of purely mesenchymal origin, we asked to what extent the models further resemble the tumors they were derived from besides the retained hallmark protein expression levels and patterns described above. To this end, whole-genome sequencing (WGS) and bulk mRNA sequencing (RNA-seq) of the tumor and tumoroid model at the time of drug screening (i.e., successful establishment), were performed.

Copy number profiles were first compared between tumors and tumoroid models showing that profiles are indeed highly concordant (Fig. 3A). The detected copy number alterations included those commonly observed in RMS, with genomic gains in chromosome 8 in FN-RMS and gains in chromosome 1 and 12 in FP-RMS (Shern et al., 2014; Weber-Hall et al., 1996). In addition, copy number profiles of individual RMS tumors and derived tumoroid models show a high concordance (Suppl. Fig. 2A and B). This indicates that the established models resemble the tumors they were derived from on a more global genomic level.

**Fig. 3.**
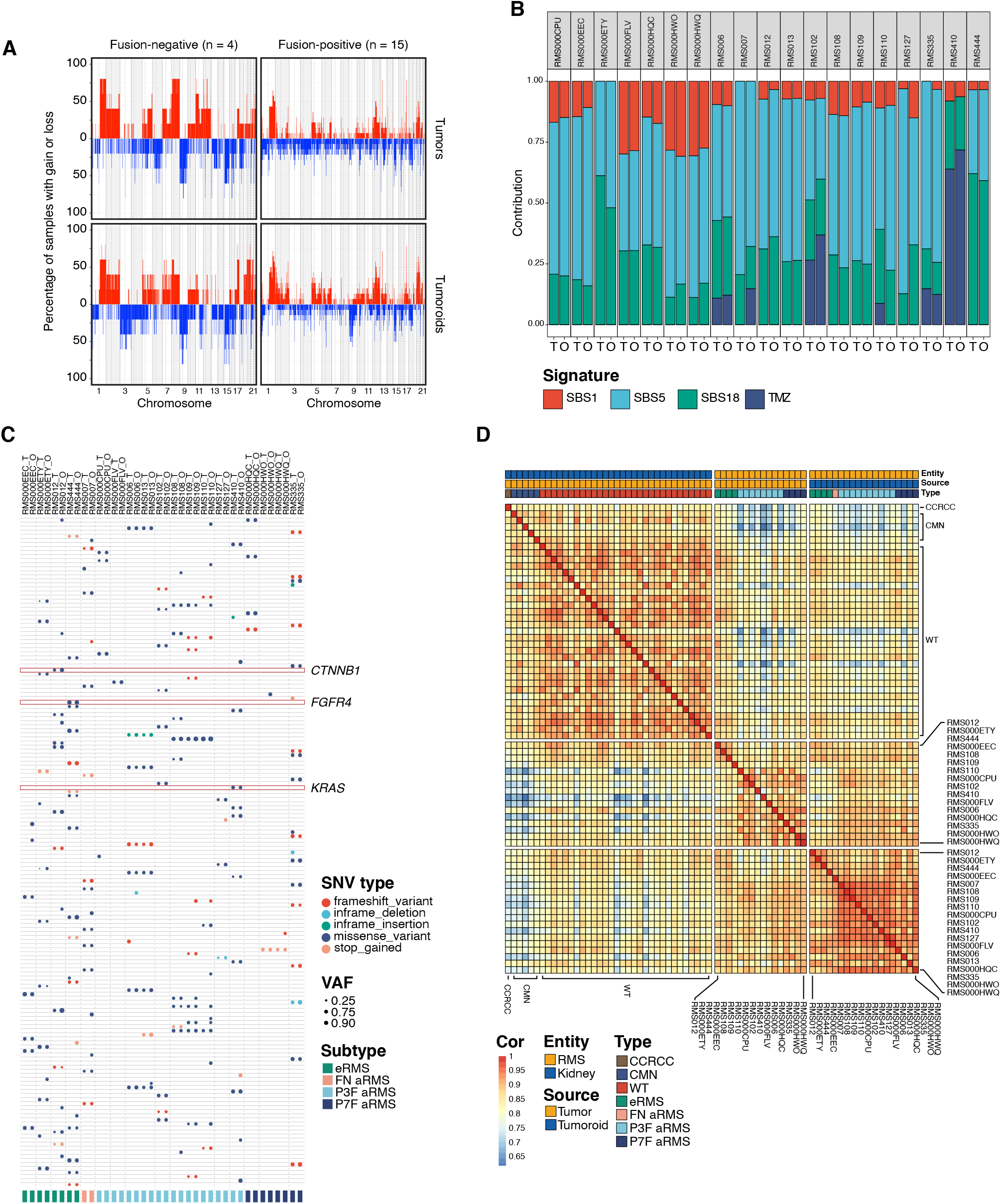
RMS tumoroid models molecularly resemble the tumor they are derived from: (A) Copy number frequency plots of RMS tumors (upper row) and derived RMS tumoroid models (lower row) divided by fusion-status (columns). Chromosomes are annotated on the x-axis from left to right while the y-axis shows the percentage of samples in this group carrying a gain (red) or loss (blue) in this genomic region. (B) Contribution of somatic mutational signatures per tumor and tumoroid model. SBS = single base substitution, TMZ = temozolomide, T = tumor, O = tumoroid. (C) Table depicting pathogenic single-nucleotide variants (SNVs) in RMS tumors (T) and tumoroid models (O). Circle color indicates SNV type while circle size indicates variant allele fraction (VAF). Vertical dotted lines separate samples derived from individual patients. Highlighted are genes previously reported for this RMS subtype. (D) Correlogram of bulk mRNA sequencing expression profiles of pediatric kidney tumors (controls) as well as RMS tumoroid models and RMS tumors. CCRCC = Clear Cell Renal Cell Carcinoma, CMN = Congenital Mesoblastic Nephroma, WT = Wilms Tumor, Cor = correlation.

Various mutational processes are active in cells, which cause distinct somatic mutational signatures. These signatures are characterized by specific patterns of single base substitutions (SBS) in the context of their two flanking bases (Alexandrov et al., 2020). The presence of certain somatic mutational signatures in a cell can be associated with the underlying mutational processes. These processes are not restricted to *in vivo* systems, but can also occur during culture (Petljak et al., 2019), forming a potential source of genomic destabilization. To test whether the somatic mutational signatures and thus the underlying mutational processes present in the tumors (“T”) are concordant with those in the derived tumoroid models (“O”), we first measured the relative contributions of different signatures per sample. The main signatures observed are signatures associated with aging (SBS1 and SBS5), as well as a signature associated with increased oxidative stress (SBS18) (Alexandrov et al., 2020), and a signature associated with exposure to the chemotherapeutic Temozolomide (TMZ) (Kucab et al., 2019) used in the treatment of RMS (Defachelles et al., 2021) (Fig. 3B). Importantly, the signatures detected in the tumoroid models are highly concordant with those detected in the original tumors (Fig. 3B). In line with this, no global differences in SBS profiles between RMS tumors and derived tumoroid models are detected when analyzed collectively (Suppl. Fig. 2C). Lastly, we calculated the similarities of the individual SBS profiles of all tested tumor and tumoroid model samples. Tumoroid models cluster with the tumors they are derived from, indicating that the mutational landscape is retained in the models (Suppl. Fig. 2D). Only the tumor and tumoroid model of RMS127 do not cluster closely, likely due to derivation from a bone marrow aspirate with low tumor cell infiltration (5-10% as estimated by pathology). Overall, there is a high concordance between the somatic mutational signatures of RMS tumors and the derived tumoroid models, again illustrating that the established models resemble the patient tumors.

While FP-RMS are driven by the prototypical fusion genes, FN-RMS are characterized by SNVs in known oncogenes such as *TP53* or *RAS* family members (Shern et al., 2014). Currently available preclinical models of RMS do not cover the full spectrum of these SNVs, thus limiting the applicability to test novel targeted drugs in RMS. To investigate whether the established RMS tumoroid models harbor any of these SNVs and whether these were already present in the tumor, all SNVs predicted to be pathogenic for protein function and with a variant allele fraction (VAF) of above 0.3 in either tumor and/or tumoroid were evaluated. As already indicated by the mutational signature analyses, RMS tumoroid models retain SNVs present in the tumor to a high degree (Fig. 3C). Similarly, samples from the same patient but from different sites or acquired from different relapses also show a high overlap in SNVs (Fig. 3C).

Two of the FN-eRMS tumoroid models harbor previously described oncogenic mutations in FN-eRMS, i.e., *CTNNB1* (p.T41A) in RMS012 and *FGFR4* (p.V550L) in RMS444 (Shern et al., 2014). To our knowledge, these are the first preclinical RMS models harboring these specific mutations. Moreover, one *PAX3-FOXO1* FP-aRMS (RMS410) displays an oncogenic mutation in *KRAS* (p.G12A), which is uncommon in FP-RMS (Shern et al., 2014). Lastly, two FN-RMS tumoroid models (RMS007 and RMS012) display non-annotated frameshift mutations in the *BCOR* gene with a high VAF of above 0.9. Mutations in *BCOR* have been reported in RMS with a higher prevalence in FN-RMS compared to FP-RMS (Shern et al., 2014). The mutation analysis shows that RMS tumoroid models not only retain specific SNVs already present in the tumor but that these models also contain mutations for which no preclinical model was previously available.

mRNA expression profiles of FN- and FP-RMS are fundamentally different, mainly due to the transcriptome-wide impact of the fusion transcript in FP-RMS (Wachtel et al., 2004). Additionally, the transcriptional program of *in vitro* cultured organoids is influenced by the culture conditions and can deviate from the transcriptional program of the parent tissue (Lu et al., 2021). Given that FN-RMS and FP-RMS tumoroid models are cultured in the same medium, we asked whether the transcriptional differences observed between the original tumors are retained in the models. Analysis of RNA-seq shows a high correlation between the expression profiles of RMS tumoroid models with the same fusion status (i.e., FP-RMS vs. FN-RMS, Fig. 3D), as has previously been shown for primary RMS tumor samples (Wachtel et al., 2004). Furthermore, correlation between the expression profiles of RMS tumor and tumoroid models of the same fusion-type is high, while correlation with control kidney tumors is low, showing that the fundamental differences in the expression profiles of FN- and FP-RMS are retained in culture (Fig. 3D). In summary, the early tumor cell detection by RT-qPCR, the morphological and marker protein analyses, WGS for copy number profiles, for somatic mutational signatures and for individual SNVs, as well as the comparative transcriptomic analyses, indicate that the RMS tumoroid models resemble the original RMS tumor they are derived from.

### Genetic and transcriptional stability of tumoroid models over time

We next asked whether the models remain genetically and transcriptionally stable after culture over extended periods. Although convenient for certain experimental approaches, conventional RMS cell lines can acquire new mutations through culturing. An example is the widely used *PAX3-FOXO1* FP-aRMS cell line RH30, that harbors a *TP53* mutation, uncommon in primary FP-RMS (Felix et al., 1992; Shern et al., 2014). To investigate stability, a subset of RMS tumoroid models were kept in culture over three to six months until they reached passage 40 (“OL” for late passage, as compared to “O”, the standard passage analyzed here and sufficient for drug screening) and characterized again by WGS and RNA-seq. Furthermore, two independently derived tumoroid models (i.e., established from the same tumor piece of which a part was cryo-preserved after the first tumoroid establishment) were included in this comparison (“O2”) to assess the robustness of the establishment protocol.

RMS tumoroid models show long-term propagation potential with five out of seven lines tested reaching passage 40 and two lines dropping out at passage 17 and 20, respectively. As drug screening is usually performed between passage 6 to 12, this shows that models can be readily used after such an initial screening. Comparison of individual copy number profiles of standard and late passage (Suppl. Fig. 3A), as well as independently derived paired tumoroid models (Suppl. Fig. 3B) shows no major copy number differences between the respective models. Furthermore, analysis of somatic mutational signatures shows that the contributions of identified somatic mutational signatures (Fig. 4A) as well as the contributions of individual SBSs between models (Suppl. Fig. 3C) are highly similar. Lastly, analysis based on SBS profiles shows clustering of models derived from individual patients (Suppl. Fig. 3D). The exception to this is RMS335 “OL” which may be due to a mutation in the DNA damage response gene *ATR* in both the tumor and the tumoroid sample, potentially resulting in the gain of new mutations during culturing. The similarity between SBS profiles of RMS335 “O” and “OL” was nevertheless high. Taken together, these analyses show that RMS tumoroid models generally remain stable, even over extended periods of culturing and that the establishment protocol is robust, yielding highly similar models when independently derived from a single tumor sample.

**Fig. 4.**
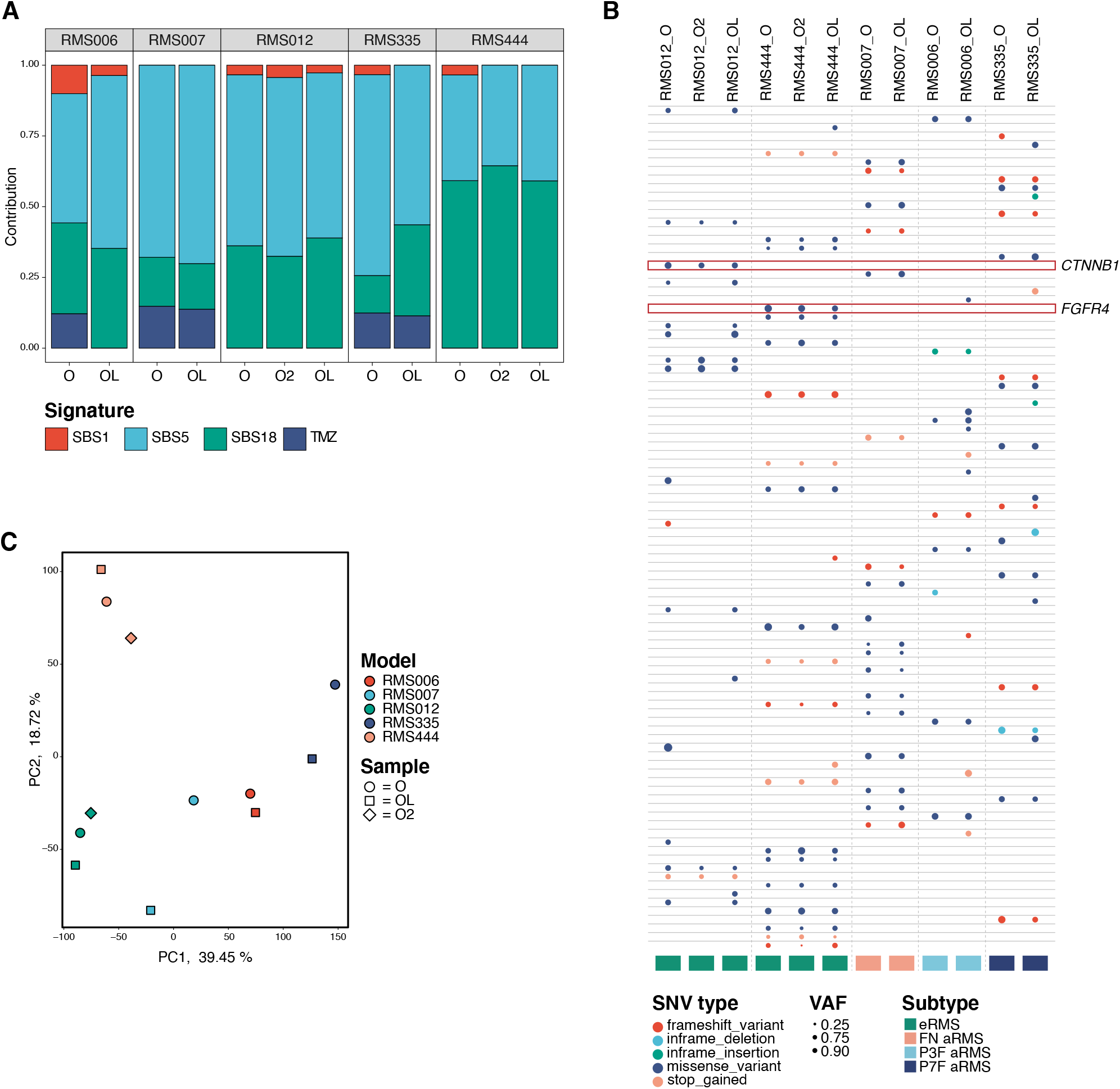
Genetic and transcriptional stability of tumoroid models over time: (A) Contribution of somatic mutational signatures per tumor and tumoroid model. SBS = single base substitution, TMZ = temozolomide, T = tumor, O = standard passage tumoroid model, OL = late passage tumoroid model, O2 = independently derived tumoroid model. (B) Table depicting pathogenic single-nucleotide variants (SNVs) in different RMS tumoroid models derived from the same tumor sample. Circle color indicates SNV type while circle size indicates variant allele fraction (VAF). Highlighted are genes previously reported for this RMS subtype. (C) Principal component analysis on bulk mRNA sequencing derived global gene expression. Color indicates the RMS tumoroid model while the symbol indicates the sample. O = standard passage tumoroid model, OL = late passage tumoroid model, O2 = independently derived tumoroid model.

As before (see Fig. 3C), individual pathogenic SNVs were also evaluated. Importantly, the majority of SNVs are retained after long-term culturing, with the models acquiring only a few additional SNVs (Fig. 4B). In addition, the models independently derived from the same tumor show a high overlap of pathogenic SNVs (Fig. 4B). Notably, the afore-mentioned oncogenic mutations in *CTNNB1* and *FGFR4* are detected in all related samples (Fig. 4B). This further indicates that the models do not lose key mutations and that the establishment protocol results in the outgrowth of representative tumor cells harboring these mutations. RNA-seq expression profiles from the standard passage, late passage, as well as the independently derived tumoroid models were compared to test whether the core transcriptional program of RMS is retained. Principal component analysis shows that global expression profiles of tumoroid models derived from the same patient cluster together (Fig. 4C). This suggests that our models are not only genetically, but also transcriptionally stable over time.

### RMS tumoroid drug screening reflects established drug sensitivities

Having established that the RMS tumoroid models resemble the tumors they are derived from with stability during prolonged culturing, their suitability for research was further investigated in two specific ways, i.e., via drug screening and CRISPR/Cas9 genome editing. Development of novel RMS treatments has been hampered by a lack of preclinical models that can efficiently be subjected to drug screening. To be of use for such screening approaches, it is imperative that new models reflect drug sensitivities already known for these tumors. This was investigated with a custom pediatric cancer library of 165 compounds, comprising standard of care chemotherapeutics as well as early-phase clinical trial targeted compounds. To mimic the three-dimensional configuration of tumors, cell plating was optimized so that tumoroid cells form homogenous 3D spheres in 384-well plates. The protocol also included prior growth curve determination of the number of cells that must be plated for each individual tumoroid model to prevent overgrowth during screening (Materials & Methods).

Vincristine and actinomycin D are routinely used in RMS treatment and indeed show broad efficacy in all models tested (Fig. 5A, bottom highlighted box). Furthermore, the proteasome inhibitor bortezomib shows a similarly high efficacy in all tested models. This is in line with previous studies showing that both major histological RMS subtypes are susceptible to bortezomib treatment *in vitro* (Bersani et al., 2008) and *in vivo* (Manzella et al., 2020), suggesting that RMS tumoroid models indeed reflect drug sensitivities known for RMS tumors.

**Fig. 5.**
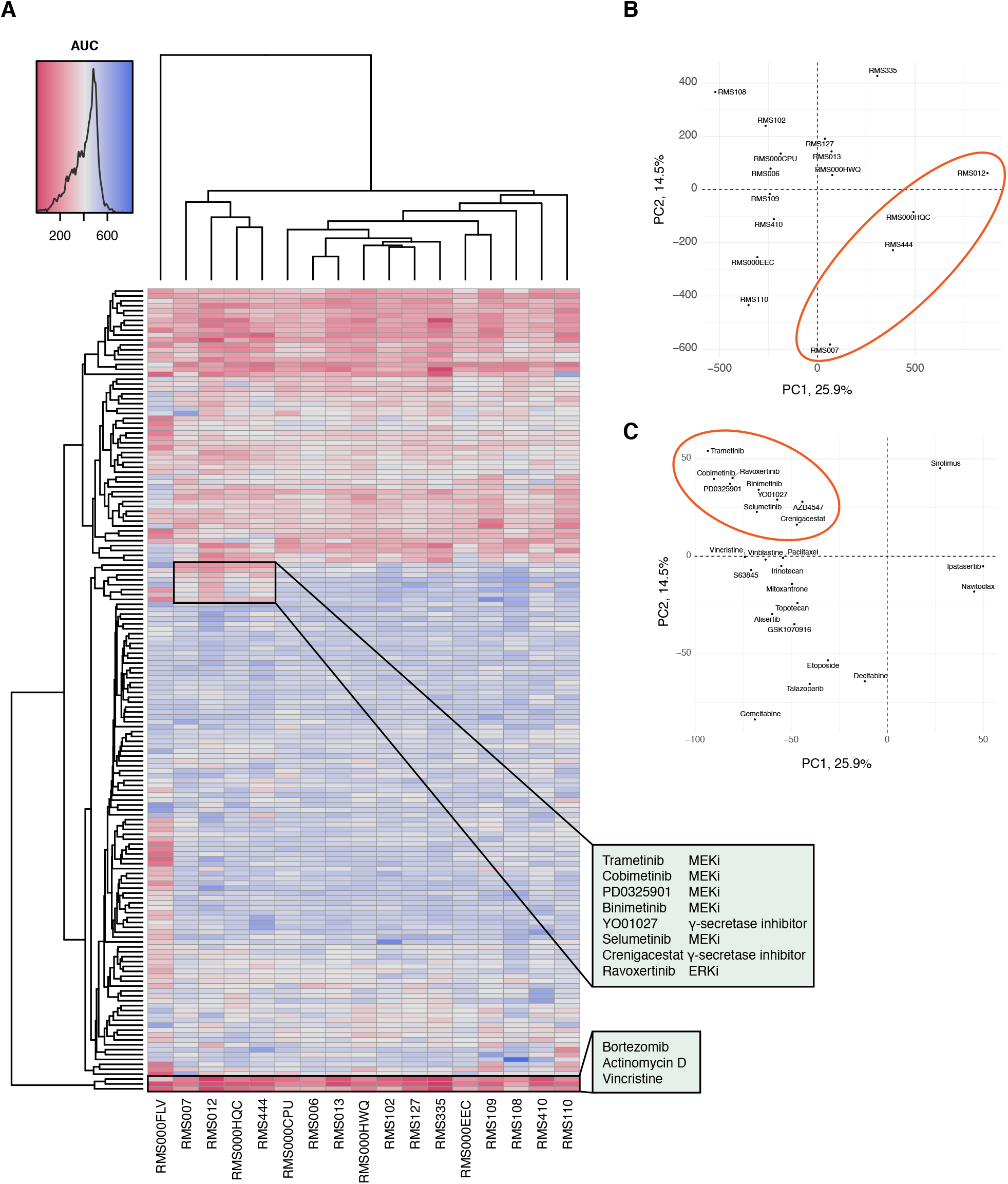
RMS tumoroid drug screening reflects established drug sensitivities: (A) Clustered heatmap of viability measurements per RMS tumoroid model (x axis) and drug (y axis), showing the Area Under the Curve (AUC) after treatment of the cells for 120 h with a dose-range of 0.1 nM to 10 µM. Low AUC (red) indicates high drug efficacy while high AUC (blue) indicates low drug efficacy. Annotated clusters of (1) MEK/ERK and γ-secretase inhibitors showing specific efficacy in RMS tumoroid models without (RMS007, RMS012, RMS444) or low (RMS000HQC) fusion transcript expression, and (2) drugs that show broad efficacy across RMS tumoroid models. (B) Principal component analysis of drug screening AUC values of the RMS tumoroid models (RMS000FLV omitted due to outlier behavior as discussed in the main text). The ellipse indicates the cluster that shows specific sensitivity against MEK/ERK and γ-secretase inhibitors. (C) Principal component analysis of the top 25 contributing drugs that influence variance. The ellipse indicates the MEK/ERK and γ-secretase inhibitors as well as AZD4547 (RMS000FLV omitted as described above).

Clustering of RMS tumoroid models based on drug efficacy shows two main groups, comprising 12 and 4 models, respectively, with one unclustered model (RMS000FLV). This model nevertheless shows high sensitivity to the afore-mentioned drugs (bottom of Fig. 5A). The outlier behavior is not caused by differences in growth during the experiment (see Suppl. Table 2) and may be explained by the fact that this is the only treatment-naïve FP-aRMS model in the collection (Fig. 1B). The group of 4 models, that contains all successfully screened FN-RMS tumoroid models, is more sensitive to all tested inhibitors of MEK/ERK (MAPK signaling pathway) as well as the two inhibitors of γ-secretase (NOTCH signaling pathway) in the drug panel, when compared to the other group containing only FP-RMS tumoroid models (Fig. 5A, top highlighted box, and Fig. 5B and C). Importantly, sensitivity of FN-RMS against inhibitors of MAPK and NOTCH signaling has previously been reported (Belyea et al., 2011; Yohe et al., 2018). This group also contains the FP-aRMS tumoroid model RMS000HQC which shows a very low expression of its fusion transcript (Fig. 2B), which potentially resulted in it displaying sensitivities more commonly observed in FN-RMS. Taken together, our results indicate that drug sensitivities observed in RMS tumoroid models reflect those known in primary RMS tumors. This shows the potential these models hold for testing novel drugs. As drug screening could be performed as early as 27 days after sample acquisition, with a median time to drug screening of 81 days, this highlights their relevance for personalized approaches.

### RMS tumoroid models can be molecularly edited using CRISPR/Cas9

The applicability of preclinical models would be further enhanced by the possibility of genetic modification. To test this, we used CRISPR/Cas9 (Jinek et al., 2012) to knock out the well-known tumor suppressor gene *TP53*. This choice is based on the recent report that loss of functional P53 protein confers a worse prognosis in RMS (Shern et al., 2021). RMS012 FN-eRMS tumoroid cells, with wildtype *TP53* as determined by WGS, were transfected with a plasmid harboring a *TP53* targeting sgRNA as well as Cas9. Successfully edited cells were selected with nutlin-3 (Drost et al., 2015) (Fig. 6A), resulting in cells with complete loss of P53 protein as confirmed by Western Blot (Fig. 6B). Sanger sequencing shows a spectrum of Indels consistent with a polyclonal population of P53 deficient cells (Suppl. Fig. 4A).

**Fig. 6.**
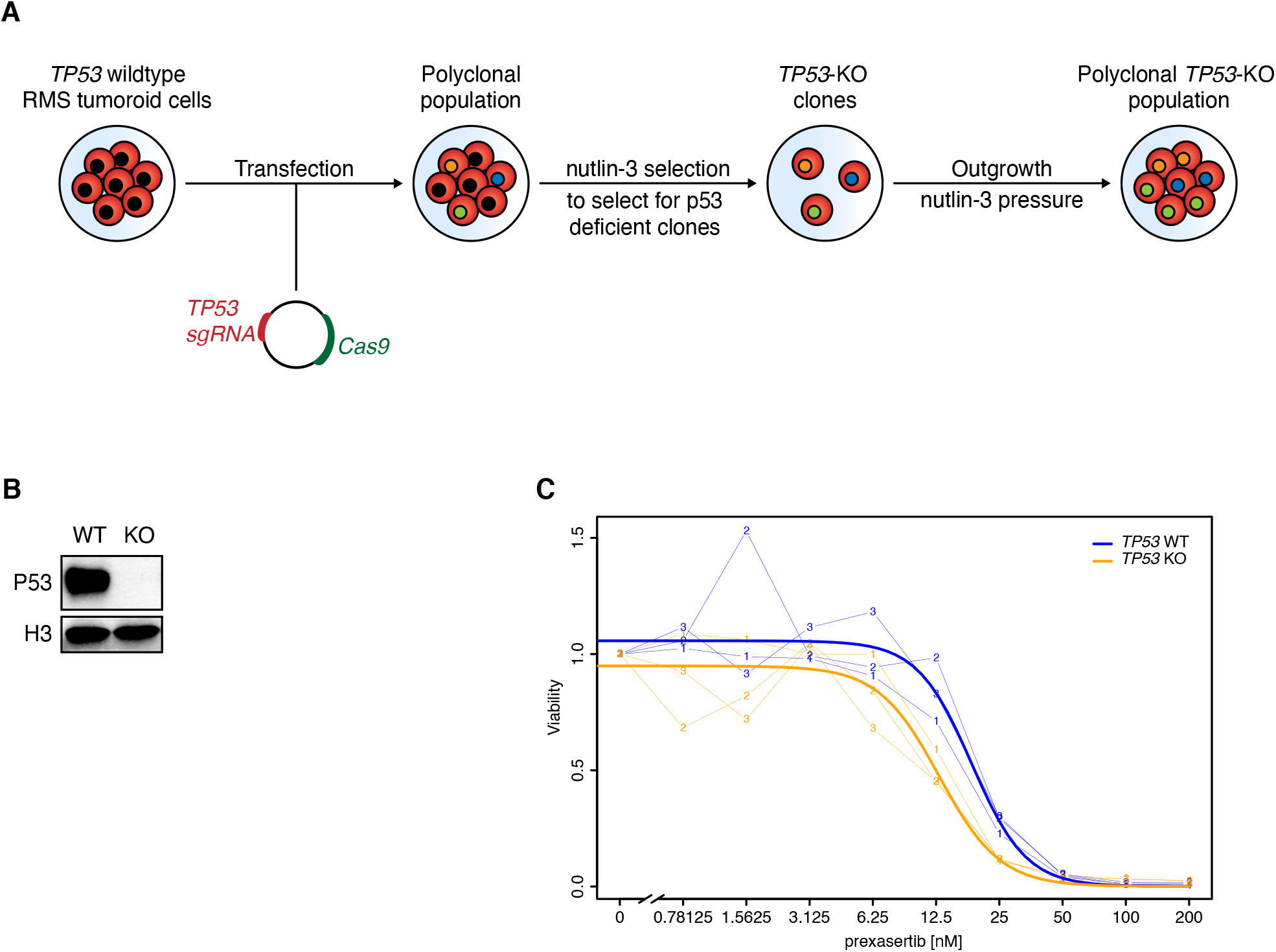
RMS tumoroid models can be molecularly edited using CRISPR/Cas9 with P53 deficient eRMS being more sensitive to the checkpoint kinase inhibitor prexasertib: (A) Transfection and selection strategy to achieve *TP53* knockout in a *TP53* wildtype tumoroid model. (B) Western Blot analysis of *TP53* wildtype (WT) and knockout (KO) RMS tumoroid line RMS012. Histone 3 (H3) served as loading control. (C) Dose-response curve of *TP53* WT and KO cells treated with the Chk-1 inhibitor prexasertib. Thin lines with numbers indicate individual technical replicates while thick lines indicate fitted lines over all replicates.

### P53 deficient eRMS tumoroid cells are more sensitive to the checkpoint kinase inhibitor prexasertib

In light of the recently reported prognostic significance of P53 loss in RMS (Shern et al., 2021), we investigated what implications loss of P53 might have. P53 governs various cellular functions, including response to DNA damage through control of the G1/S checkpoint during cell cycle progression by halting the proliferative machinery to give cells time to repair DNA damage (Yonish-Rouach et al., 1991). The rate of DNA double-strand breaks (DSBs) was therefore first examined in the P53 deficient RMS tumoroid cells by measuring phosphorylation of histone H2AX serine 139, a proxy for DNA DSBs (Rogakou et al., 1998). No difference in the amount of DNA DSBs was observed between *TP53* wildtype and knockout cells however (Suppl. Fig. 4B). To avoid catastrophic failure of the proliferative machinery and avoid cell death, P53 deficient cells are dependent on the G2/M checkpoint to repair DNA DSBs (Dobbelstein and Sørensen, 2015). Inhibitors of the checkpoint kinase 1 (Chk1), which controls the G2/M checkpoint, have therefore been suggested as a treatment for P53 deficient tumors, such as in certain ovarian and breast cancer subtypes (Lee et al., 2018; Ma et al., 2012). This hypothesis was tested, resulting in the observation that *TP53* knockout RMS tumoroid cells are significantly (*p* = 0.008) more sensitive to the Chk1 inhibitor prexasertib, compared to their wildtype counterpart (Fig. 6C). The absolute difference between IC_50_ values is not extreme (18.5 nM for *TP53* wildtype versus 12.9 nM for *TP53* knockout cells), likely reflecting the fact that RMS012 *TP53* wildtype cells are already sensitive to prexasertib. These results indicate that exploiting replicative stress in P53 deficient RMS is indeed an interesting therapeutic avenue to pursue. Moreover, the results demonstrate that it is possible to perform gene editing in these novel RMS tumoroid models, thereby further increasing their utility.

## Discussion

### A tumoroid collection of purely mesenchymal origin

To date, organoid technology has primarily been employed to generate models of malignant tumors of epithelial origin (i.e., carcinomas) (Bleijs et al., 2019). Feasibility to use this technology on non-epithelial cancer has only recently been shown (Abdullah et al., 2021; Fusco et al., 2019; Jacob et al., 2020; Saltsman et al., 2020; Yamazaki et al., 2021). Here we extend the tumor organoid approach, demonstrating applicability to tumors of entirely mesenchymal origin (i.e., sarcomas), resulting in only the second thoroughly characterized tumoroid collection specific for pediatric cancer.

Two factors likely contributed to the delayed adaptation of organoid technology to sarcomas. First, the technology was developed for healthy epithelial tissue, followed later by translation to the corresponding cancer entity, as in the case of colorectal carcinoma (Sato et al., 2009, 2011; Van De Wetering et al., 2015). The cell of origin of RMS is still not fully characterized (Hettmer and Wagers, 2010). Therefore, extensive culture optimization of healthy tissue first, with translation to cancerous tissue later, has not been feasible for RMS. Consequently, optimization had to be conducted on tumor samples, which are not readily available. Secondly, and related to the issue of tissue availability, sarcomas are far less common than carcinomas, accounting for less than 1% of all solid adult malignancies (Burningham et al., 2012). In pediatric cancer however, sarcomas are much more common, encompassing 21% of all solid tumors in children (Burningham et al., 2012). This further highlights the importance of the currently described approach and collection.

### Representation of a broad spectrum of RMS subtypes

The tumoroid models include representatives of both major histological subtypes, all major fusion types, different age groups, both sexes, treatment-naïve as well as pre-treated, primary as well as metastatic tumors (Fig. 1B and Suppl. Table 1). Rarer subtypes such as sclerosing/spindle-cell RMS (Rudzinski et al., 2015) will be exciting to include, as also indicated by a recent case report (Acanda De La Rocha et al., 2021). Compared to patient incidence rates (Glosli et al., 2021), the collection has an underrepresentation of head and neck RMS. Although RMS tumor samples from this region were acquired, models from such samples failed, regardless of subtype, clinical stage, sample quantity or quality. Interestingly, this indicates that RMS arising in the head and neck may depend on specific factors that have not yet been identified. Studies in genetically engineered mice indicate that aberrant Hedgehog signaling can give rise to FN-RMS from non-myogenic endothelial progenitors in the head and neck (Drummond et al., 2018). Here, activation of Hedgehog signaling by Smoothened agonists did not facilitate establishment of head and neck RMS tumoroid models, indicating that additional factors may be necessary. Regardless of such future developments, the protocol described here yields models that can be rapidly established from a broad range of quite different RMS subtypes.

### New models for RMS research

Currently available preclinical models of RMS include conventional cell lines, various genetically engineered animal models (GEMs), as well as patient-derived xenograft models (PDX or O-PDX when transplanted orthotopically) (Imle et al., 2021; Kashi et al., 2015). Each of these systems possesses specific benefits and drawbacks, resulting in suitabilities for different research questions or stages in drug development (Kim et al., 2020). While conventional cell lines are easy to use, with low costs, large-scale screening potential and ease of genetic modification, due to prolonged culturing they often do not recapitulate many basic features of the genetic and molecular background of the tumor they were derived from, thus possessing only limited predictive value (Hinson et al., 2013). GEMs, on the other hand, are well-suited for cell of origin studies and can provide valuable insights into cancer onset mechanisms. GEMs are usually not suitable for high-throughput screening, mostly due to the low tumor penetrance or the intricacies of animal studies (Kersten et al., 2017). In contrast to GEMs, in O-PDX models, tumor samples are transplanted onto immune-deficient mice to allow for engraftment, growth and later propagation of the tumor tissue from mouse to mouse. Tumors propagated in this manner are thought to be genetically stable over time and to reflect the patient tumor they were derived from, giving them predictive value concerning preclinical drug testing (Gao et al., 2015). Disadvantages include the necessary use of mice as hosts and the tumor-entity specific engraftment time which can take up to several months (Pompili et al., 2016). Furthermore, there is evidence that not all PDX models are genetically stable (Petljak et al., 2019).

The RMS tumoroid models combine several of the above-mentioned benefits. They can be rapidly established and expanded like conventional cell lines, enabling drug screening. As with cell lines, molecular editing to mimic certain disease backgrounds is possible using CRISPR/Cas9. Unlike cell lines however, tumoroid models depend on defined media including recombinant growth factors as well as an ECM substitute, resulting in higher costs. RMS tumoroid models molecularly resemble the patient tumor they are derived from, sharing this characteristic with O-PDX models, while possessing the above-mentioned advantages of rapid establishment and expansion. Compared to O-PDX models, establishment success is lower in RMS tumoroid models, indicating that niche factors in the host mice are important for facilitating establishment. This may be especially important in particular subtypes such as RMS from the head and neck region. On the other hand, RMS tumoroid models are less intricate in their propagation. Lastly, while GEMs are considered essential for cell of origin studies, recent advances have shown that by genetic editing, such studies can now also be performed in tumoroid models (Custers et al., 2021).

In conclusion, we established a well-characterized, well-annotated collection of RMS tumoroid models, being the first such collection of tumoroid models derived from purely mesenchymal malignant tumors (i.e., sarcomas) and only the second comprehensive tumoroid model collection derived from pediatric cancer (Calandrini et al., 2020). This collection contains all major subtypes of RMS and the models can be used for drug screening as well as molecular editing. An interactive, browser-based companion Shiny app (https://rmsdatabiobank.shinyapps.io/app1_3/) that makes all the described data easily accessible, accompanies this paper. The RMS tumoroid models will be a useful complementary system to study the biology of RMS and to improve treatment.

## Acknowledgements

We thank the patients and their parents for contributing by consenting to this study; the nurses, doctors and other health care professionals in our institute with whom we work closely; our center’s high-throughput screening facility, in particular Bianca Koopmans, Kimberley Ober, and Sander van Hooff; Philip Lijnzaad, Thanasis Margaritis and Tito Candelli, as well as past and present members of the Holstege group.

## Financial support

M.M. received financial support from the Deutsche Forschungsgemeinschaft (#408083583). C.C. was supported by Foundation Children Cancer Free (#292). J.D. received support from the European Research Council (ERC) starting grant 850571 and the Dutch Cancer Society (KWF)/Alpe d’HuZes Bas Mulder award (#10218). We are grateful for the financial support provided by the Foundation Children Cancer Free (KiKa core funding).

## Author contributions

Conceptualization, M.M. and F.H.; Methodology, M.M., M.G.K. and W.B.; Investigation: M.M., M.G.K, T.d.S., W.B., E.F-M., M.B., J.D., C.C. and F.M., M.v.T.; Resources, M.D., S.E., K.L., R.K., S.T.v.S., L.H., U.F., J.M., M.v.N., B.T., J.H-K., P.K., J.M., M.v.d.W., R.v.B. and J.D..; Data curation, M.G.K., T.d.S., A.J. and H.K.; Writing – Original Draft, M.M. and F.H.; Visualization, M.M., T.d.S. and F.M.; Funding Acquisition, F.H., Supervision, M.M. and F.H.

## Conflict of interest disclosure statement

The authors declare no competing interests.

## Materials & Methods

### Tumor sample acquisition

Tumor samples of RMS were obtained via an established tumor sample acquisition route from patients treated at the Emma Children’s Hospital Amsterdam (Amsterdam UMC) (RMS006, RMS007, RMS013) or as part of the biobank initiative of the Princess Máxima Center for Pediatric Oncology, Utrecht, Netherlands (PMC) (remaining tumor samples). Ethics approval was granted for the biobanking initiative, and the PMC biobank committee granted approval for the present project. All patients and/or their legal representatives signed informed consent to have tumor samples taken for biobank usage.

A subset of patients was furthermore enrolled in a local personalized medicine trial (i.e., iTHER study) through which a subset of DNA and RNA specimens from RMS tumors were obtained.

### Tumor sample preparation for establishment of RMS tumoroid models

Solid tumor samples (i.e., needle biopsies or resection specimens) were transferred to collection medium (see below) to retain viability of cells. After pathological examination, suitable samples (i.e., containing tumor cells) were processed in a sterile work environment as follows: the sample was transferred to a sterile dish and covered with a droplet of BM1* culture medium (see below) before being minced to fine pieces using scalpels. If the tumor sample was of sufficient size, a portion of this minced mixture was stored viably (see below). The remainder of the mixture was put on a pre-wet 70 µM strainer, scrapped with a cell scraper to obtain a single-cell suspension and collected in a tube (A, single-cell fraction, cultured in 2D). Tissue fragments left on the strainer were collected in another tube (B, strainer fraction, cultured in 3D).

A: The single-cell fraction was now spun down (300 g, 5 min, 4 °C) and the supernatant was removed from the resulting pellet. If the pellet was of grey color (i.e., not containing a high percentage of red blood cells, RBCs), the pellet was resuspended in 1 ml of BM1* medium. If the pellet contained a high percentage of RBCs (i.e., by displaying a red stain), the cell pellet was resuspended in red blood cell lysis buffer (Roche) and incubated at room temperature for 5 min. Thereafter, the reaction was stopped by adding collection medium and the mixture was spun down again (as above). Again, supernatant was removed and now the pellet was resuspended in 1 ml of BM1* medium (without Basement Membrane Extract, BME, see below). In each case, cells were now counted using a TC20 Automated Cell Counter (BioRad) to get a rough estimate on viability and cell numbers. Cells were plated with a sufficient density (at least 10.000 viable cells per 1 cm^2^ of surface area), supplemented with 0.3 to 0.5 % cold BME type 2 (see below).

B: The strainer fraction was dissolved in pure cold BME (roughly one-third volume strainer fraction and two-third volume BME) and mixed thoroughly. Droplets of 5 to 10µl of this mixture were formed on 24- or 48-well pre-warmed cell culture plates and incubated for 5 min at room temperature to allow the BME to solidify. Thereafter, the cell culture plates were incubated upside-down for another 25 min at 37 °C to allow the material to “sink” to the top of the droplet. Upon completed solidification, BM1* medium (at room temperature (RT) and without BME) was added to the wells so that droplets were very fully submerged in medium.

In the case of bone marrow aspirates as tumor samples (RMS410 and RMS127), sample tubes were spun down (300 g, 5 min, 4 °C, slow break) to separate RBCs and nucleated cells (white clot at the bottom of the tube). In the case of RMS410, the normal hematopoietic system was almost entirely superseded by infiltrating tumor cells so that the clot consisted mainly of tumor cells which could directly be plated (in BM1* with BME) which resulted in rapid outgrowth of the model. In the case of RMS127, the percentage of infiltrating tumor cells was estimated by pathology to be low at circa 5 to 10%. Therefore, initial cultures (plated in BM1* and BME) from the white clot also contained normal nucleated bone marrow cells which, however, were eventually outcompeted by the rapidly growing tumor cells which overtook the culture.

### Tumoroid model culturing and propagation

Growing RMS tumoroid models were inspected regularly and showed adequate growth behavior under conventional cell culture conditions (i.e., 37 °C, 5 % CO_2_). All models were regularly tested negative for mycoplasma contamination. Estimated division times ranged from 24 to 72 hours for most models. Models were passaged once or twice per week at a confluency of 70 to 80%. For passaging, old culture medium was aspirated and cells were briefly washed with sterile DPBS (Gibco, cat no. 14190144). Cell detachment was performed using TrypLE Express Enzyme (1X, phenol red, Gibco, cat no. 12605010). Depending on the model and the BME percentage, detachment took between 3 and 10 minutes (higher BME concentrations resulting in longer detachment time). Cells were collected by flushing the well or flask with collection medium and the resulting mixture was spun down (300 g, 5 min, 4 °C). Thereafter, the supernatant was removed, and the pellet was resuspended in 1 ml of BM1* and cells were counted using a TC20 Automated Cell Counter (BioRad). Single-cell suspensions were mostly re-seeded at the same density as their parental/previous generation/passage.

### Cell culture media

#### Base medium (BM)

To prepare a 500 ml bottle of BM, Glutamax (5 ml, Gibco, cat no. 35050061), Penicillin/Streptomycin (10,000 U/ml, 5 ml, Gibco, cat no. 15140122), and B27 (without vitamin A, 10 ml, Gibco, cat no. 12587010) were added to a full bottle of advanced DMEM/F12 (500 ml, Gibco, cat no. 12634010). BM was stored at 4 °C and was used within two months.

#### Complete culture medium (BM1*)

To prepare the complete culture medium BM1*, 47.5 ml of the above-mentioned base medium (BM) was taken and pipetted into 50 ml tube. Thereafter, the components below were added (no specific order). BM1* was stored at 4 °C and was used within 7 to 10 days (thereafter, the stability of the growth factors may be compromised).

Components:

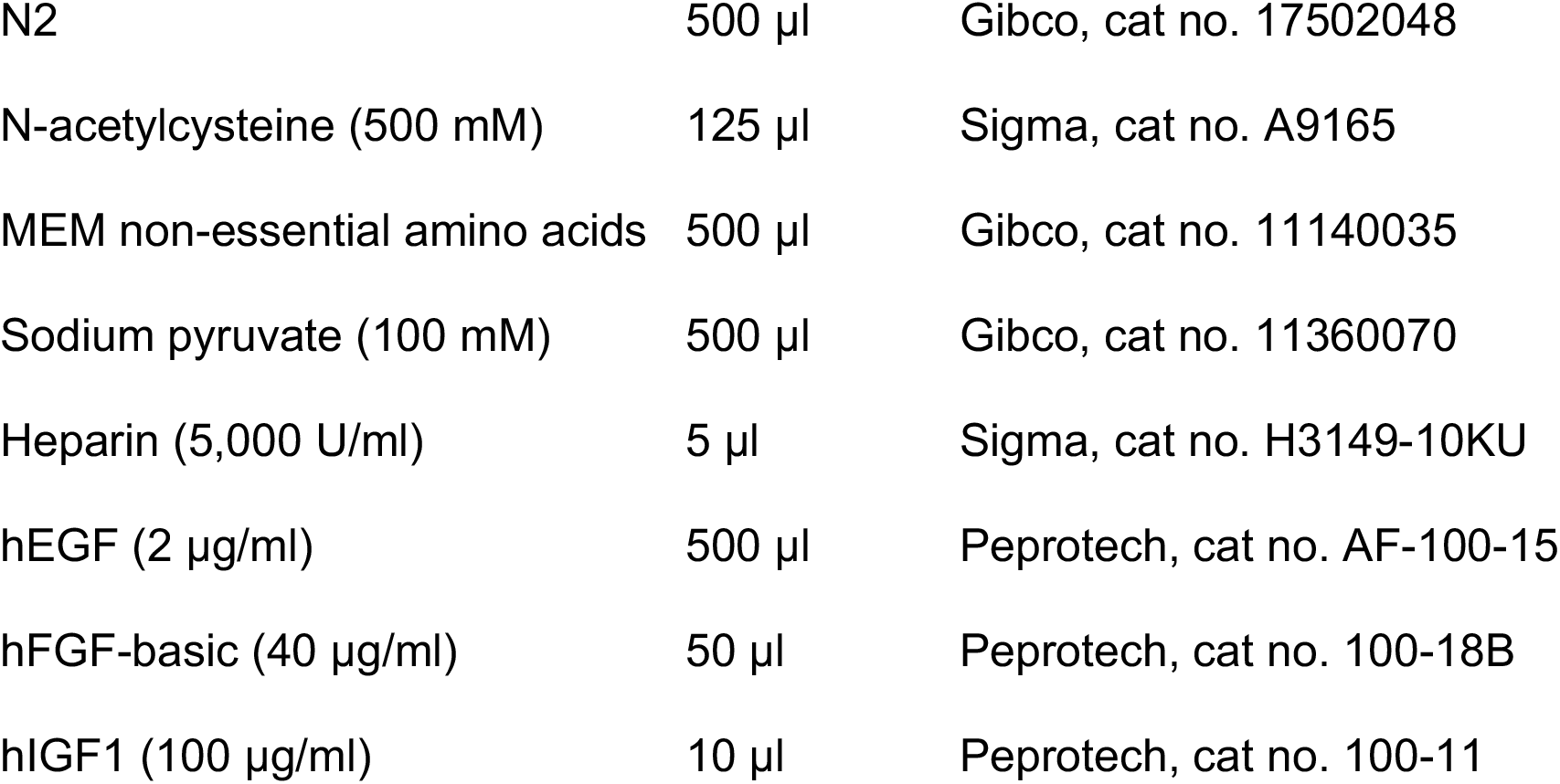
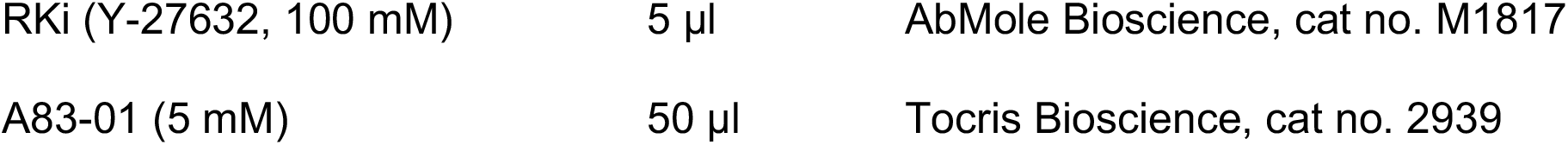

#### Collection medium

To prepare a 500 ml bottle of collection medium, Glutamax (5 ml, Gibco, cat no. 35050061), Penicillin/Streptomycin (10,000 U/ml, 5 ml, Gibco, cat no. 15140122), and HEPES (1 M, 5 ml, Gibco, cat no. 15630049) were added to a full bottle of advanced DMEM/F12 (500 ml, Gibco, cat no. 12634010). Collection medium was stored at 4 °C and was used within two months.

### Basement Membrane Extract type 2 (BME)

To facilitate attachment of cells, culture medium was supplemented with 0.1 to 0.5 % BME (Cultrex Reduced Growth Factor Basement Membrane Extract, type 2, Pathclear, R&D Systems, 3533-005-02). We observed batch-to-batch variations of this product, resulting occasionally in suboptimal attachment of cells when the BME concentration was too low. On average, 0.2 to 0.3 % BME supplementation was sufficient for stably established RMS tumoroid models. However, upon encountering suboptimal cell attachment, BME concentrations were raised to 0.5 % in established cultures. During the initial establishment process, higher concentrations of BME (0.3 % to 0.5 %) showed increased attachment rates of cells.

### Freezing and storing procedure

RMS tumor samples as well as established RMS tumoroid cultures were viably frozen as follows: a sufficient number of viable cells (at least 0.5×10E6, mostly 1×10E6) were diluted in 0.5 ml BM1* medium in a cryo tube. Then, an equal volume of freeze-mixture consisting of 80 % fetal calf serum (FCS) and 20 % DMSO was added dropwise, resulting in final concentrations of 50 % BM1* medium, 40 % FCS, and 10 % DMSO. Samples were frozen slowly using a freezing container in a minus 80 °C freezer. For long term storage, frozen vials were transferred to liquid nitrogen.

### Thawing procedure

Cryo-preserved RMS tumoroid cell suspensions were quickly defrosted in a water bath at 37 °C and then immediately dissolved in washing medium (to at least 5 ml of total volume to dilute the DMSO). Samples were then spun down (300 g, 5 min, 4 °C), supernatant was removed, and the cell pellet was resuspended in 1 ml of BM1* medium. Thereafter, cells were counted using a TC20 Automated Cell Counter (BioRad). Cell viability was on average 20 to 50% lower compared to viability at the time of freezing. Cells were then plated at a proper density in BME-supplemented BM1* medium and needed one to two weeks to recover before being stable enough for further experiments. Restarting tumoroid cultures from cryo vials was possible for all tumoroid models.

### Early tumor validation by RT-qPCR

Early during tumoroid establishment (upon first or second passaging), a portion of cells was set aside for evaluation of marker gene expression. For this, cells were spun down (500 g, 5 min, 4 °C), the supernatant was removed from the pellet, the pellet was dissolved in Trizol (10 minutes incubation at RT) and was immediately processed or snap-frozen and stored at minus 80 °C until further processing. Upon processing, first the organic and aqueous phases were separated by addition of 20 % chloroform, followed by spinning down (maximum centrifugation speed, 15 min, 4 °C). The (upper) aqueous phase was further processed using the Direct-zol RNA MiniPrep Kit (Zymo Research) according to the manufacturer’s protocol, including the recommended DNAse I treatment. Quality and quantity of isolated RNA were measured using a NanoDrop OneC (Thermo Fisher Scientific). Synthesis of cDNA from isolated RNA as well as a Universal Human Reference RNA that was used as negative control or for normalization (HREF, Stratagene/Agilent # 740000) was performed using an oligo-dT primed SuperScript III Reverse Transcriptase (Invitrogen) based reaction according to the manufacturer’s protocol. RT-qPCR was performed with the obtained cDNAs testing for expression levels of *G6PD* (forward: 5’-ACGGCAACAGATACAAGAAC-3’, reverse: 5’-CGAAGTGCATCTGGCTCC-3’; product size: 86 bp) (Amary et al., 2007), *DES* (forward: 5’-CCGTGGTCTCTTACTTTCCTTT-3’, reverse: 5’-CCCACTTTCTCTCCTTCTCAATC-3’; product size: 119 bp), *MYOG* (forward: 5’-TGCCCAACTGAGATTGTCTTC-3’, reverse: 5’-CTGCTACAGAAGTAGTGGCATC-3’, product size: 81 bp), *MYOD1* (forward: 5’-GTAGCAGGTGTAACCGTAACC-3’, reverse: 5’-CACACCATGCCTCAGAGATAAA-3’, product size: 148 bp), the *PAX-FOXO1* fusion transcript (forward: 5’-CCGACAGCAGCTCTGCCTAC-3’, reverse: 5’-TGAACTTGCTGTGTAGGGACAG-3’, product size: 171 bp for *PAX3-FOXO1* and 159 bp for *PAX7-FOXO1*) (Ponce-Castañeda et al., 2014) as well as the *PAX3-WWTR1* fusion transcript (forward: 5’-AGCACCAGGCATGGATTT-3’, reverse: 5’-TTCGAGGTCTGTGTCTAGGT-5’, product size: 192 bp). Expression levels of *DES*, *MYOG*, *MYOD1*, and *PAX-FOXO1* were normalized to *G6PD* (housekeeping gene) and referenced to the corresponding expression levels in the HREF using the ΔΔCq method (for the fusion transcript only normalization to *G6PD* as the lack of a fusion expression in HREF did not permit a further reference step).

### Immunohistochemistry (IHC) and H&E stainings

To perform IHC, tumoroid models were grown as floating 3D spheres. For this, 1-3×10E6 viable cells were put into an ultra-low attachment culture flask (Corning Ultra-Low Attachment 75 cm^2^ U-Flask, Corning, cat no. 3814) in BM1* but without BME supplementation. Establishment of spheres of sufficient size took between 7 and 12 days, depending on the growth characteristics of the respective tumoroid model. Spheres were harvested by carefully transferring the sphere-containing medium from the flask to a 15 ml tube and sedimenting on ice for 10 minutes. Thereafter, the supernatant was removed, and the sphere-containing pellet was resuspended in cold PBS to wash off any remaining medium. The mixture was again sedimented (see above), and PBS was aspirated. Spheres were now fixed using formalin 10 % (v/v), (= 4 % (w/v) HISTO GRADE, neutralized (pH 7.0 ± 0.2), J.T. Baker, 3933.9020 VWR) for 96 hours at 4 °C after carefully transferring them to a glass vial. The fixed spheres were then washed twice with PBS and dehydration was performed by adding ethanol solutions with increasing percentages (25 % EtOH for 15 min, 50 % EtOH for 15 min, 70 % EtOH for 15 min - after this step, spheres were stored at 4 °C and further processed in batches). Now, spheres were stained with 0.8 g/l Eosin Y dissolved in 96 % EtOH (Sigma, E4009) for 30 min and subsequently incubated three times with 100% EtOH for 30 min each. Thereafter, spheres were incubated in n-Butanol (three times 30 min) and melted paraffin (three times) before they were Paraffin-embedded using the HistoCore Arcadia H (Leica Biosystems) following the manufacturer’s protocol in a medium size mold. Hardened paraffin blocks were cut into 4 µm slices using a microtome (HM 355S Automatic Microtome, Thermo Scientific). Slides were further processed by deparaffinization (incubation with xylene three times for 3 min, then 100% EtOH two times for 3 min, 95% EtOH two times for 1 min, 70% EtOH once for 1 min, and 50% EtOH once for 1 min) and rehydration (tap water). Thereafter, antigen retrieval was performed by boiling samples for 20 min in with citrate buffer (pH 6, for Desmin staining) or Tris-EDTA-Tween buffer (pH 9, for Myogenin and MYOD1). After washing (in TBS-0.025% Triton, twice 5 min) and blocking (1.5 h in TBS-1%BSA), slides were incubated with primary antibody (α-Desmin 1:400, Abcam ab15200 rabbit antibody; α-Myogenin 1:400, Santa Cruz 5FD mouse antibody; α-MYOD1 1:200, Cell Marque EP212 rabbit antibody) overnight at 4 °C. The next day, slides were washed twice with PBS and incubated with secondary antibodies (Desmin and MYOD1: anti-rabbit-HRP, BioRad 170-6515; Myogenin: anti-mouse HRP, BioRad 170-6516) 1:500 in PBS-1% BSA. Stainings were visualized using Liquid DAB+ 2-component system (3,3’-diaminobenzidine, DAKO, Agilent K3467) following the manufacturer’s protocol and washed three times with TBS. Counter-staining was performed by incubation with thionine (0.05 % for 20 min). After subsequent incubation with 96 % EtOH, 100 % EtOH and xylene, slides were mounted using Permount mounting medium (Fisher Scientific SP15-100) and visualized using a Leica DMi6 microscope.

H&E stainings were performed manually (steps: xylene three times for 5 min, 100 % EtOH twice for 1 min, 95 % EtOH twice for 30 sec, 70 % EtOH for 30 sec, washed in demi-water, incubation with hematoxylin (Hematox 7211) for 2:45 min, washed with demi-water, brief incubations with acidic EtOH, washed with demi-water, 95 % EtOH for 30 sec, eosin incubation for 2:45 min, EtOH 70% for 30 sec, EtOH 95 % for 30 sec, EtOH 100 % twice for 30 sec, xylene three times for 1 min) or were performed at the in-house pathology department following standard protocols. Visualization was performed as described above.

The quality of stainings was evaluated by an in-house pathologist. Representative images of stained spheres are shown and were compared to H&E as well as IHC stainings obtained in pathology for the RMS tumor sample the tumoroid was derived from (representative images chosen by the pathologist).

### RNA and DNA isolation from tumor and derived tumoroid samples for whole-genome sequencing (WGS) and bulk mRNA sequencing (RNA-seq)

#### Tumoroid models

To isolate RNA and DNA for WGS and RNA-seq, tumoroid cells were collected as a pellet, snap frozen and stored at minus 80 °C. Isolation of RNA and DNA from the same pellet was performed using the AllPrep DNA/RNA/miRNA Universal Kit (Qiagen # 80224) according to the manufacturer’s protocol. Lysis in RLT buffer was followed by homogenization using a Qiashredder column as described. The flowthrough fraction in RLT buffer was used for the Allprep DNA and RNA isolations. Quality and quantity of isolated RNA and DNA were measured by using the NanoDrop OneC (Thermo Fisher Scientific), Bioanalyzer 2100 (Agilent), Qubit Fluorometer (Thermo Fisher Scientific), and marker-checks using RT-qPCR (as above).

#### Tumor samples and germline control samples (EDTA blood)

Resected tissue and/or biopsies were processed within 10 minutes after removal from the patient. DNA and RNA were isolated from the same piece of fresh frozen tissue using the AllPrep DNA/RNA/miRNA Universal Kit (Qiagen # 80224) using the QIAcube Connect (Qiagen). Reference DNA was isolated from peripheral white blood cells (EDTA blood) using the same method.

### Whole-genome sequencing (WGS) of tumor and tumoroid samples

#### Sequencing

150 ng of total DNA was used for library preparation using the KAPA HyperPlus kit (Roche), according to manufacturer’s instructions. Libraries from tumor and normal tissue were pooled in a 3:1 ratio, with a total of 7 tumor/normal pairs per S4 sequencing kit. Libraries were sequenced using 2×150 cycles on a NovaSeq 6000 (Illumina).

#### Pre-processing, alignment, and annotation

The WGS sequencing data were processed as per the GATK 4.0 best practices workflow for variant calling, using a wdl and cromwell based workflow. Reads were aligned to GRCh38 using bwamem (v0.7.13), and quality control (QC) was performed using FastQC (v0.11.5) and picardTools (v2.20.1). Somatic variants were identified using Mutect2 from GATK v4.1 and annotated using Vep (v92). Likewise, copy-number alterations (CNAs) were identified using GATK v4.1.

#### Identification of non-synonymous single nucleotide variants (SNVs)

Raw VCF files from germline, tumor and tumoroid samples were processed with vcftools (v0.1.13) (Danecek et al., 2011). Only variants that met the criteria (passed all quality filters in addition to PHRED quality score 100 (250 for indels) and minimum read depth of 10) were kept for further analyses. Indels were also filtered out if the minimum mapping quality (MQ) was below 60. Bcftools (https://github.com/samtools/bcftools) was used to remove all common snps (VAF ≥ 1

%) present in dbSNP (https://www.ncbi.nlm.nih.gov/snp/, v151 downloaded in August 2020). Filtered files were then loaded into *R* (v4.0.2) and processed with the package VariantAnnotation (Obenchain et al., 2014) and packages from the tidyverse. Variants in tumor and tumoroid models also present in the germline sample were removed from further analyses; remaining variants were filtered for VAFs > 0.3 and those tagged as missense variant, stop gained, stop lost, start lost, inframe insertion, inframe deletion, and frameshift variant were selected as non-synonymous somatic mutations. Figures were generated with ggplot2 (v.3.3.2).

#### Signature analysis

Somatic signature analysis was performed using the *R* programming language (v3.6.3) and the *R* packages MutationalPatterns (v3.2.0) and VariantAnnotation (v1.32.0) (Blokzijl et al., 2018; Manders et al., 2021; Obenchain et al., 2014). Somatic variants were filtered on both the variant allele fraction (VAF) and depth (DP). For both the control and the tumor/tumoroid samples we used DP ≥ 20. For the tumor/tumoroid samples we used VAF > 0.3, while in control samples the VAF had to be 0. Samples with 50 or less mutations would have been excluded, but this was not the case for any samples in our cohort. A mutation matrix was generated that shows how often each of the 96 types of base substitutions occurred in each sample. A variational Bayesian non-negative matrix factorization (NMF) was performed on this matrix to extract four *de novo* mutational signatures for base substitutions. The cosine similarities were then calculated between these *de novo* signatures and a set of signatures consisting of both the COSMIC signatures (v3.2, GRCh38) and the SIGNAL exposure signatures (Alexandrov et al., 2020; Kucab et al., 2019). The four signatures most similar to the *de novo* signatures (SBS1, SBS5, SBS18, and Temozolomide..200.uM..1 [TMZ]) were then used for signature refitting. Signature refitting was performed using the *fit_to_signatures_strict* function using the *best_subset* method with a *max_delta* of 0.004. Our approach of first performing *de novo* signature extraction followed by refitting is similar to the approach suggested by Maura *et al*. (Maura et al., 2019). To determine the similarity between the samples, the cosine similarities of their base substitution profiles were calculated. This resulted in a cosine similarity matrix that was used to calculate the distance between samples, which was then used for hierarchical clustering.

### Bulk mRNA (RNA-seq) of tumor and tumoroid samples: transcriptional profile and gene fusions

#### Sequencing

300 ng of total RNA was used for library preparation using the KAPA RNA HyperPrep kit with RiboErase (Roche), according to manufacturer’s instructions. The protocol was optimized to achieve an insert size of ∼300-400 nt.

RNA libraries were pooled with a total of 25 samples per S1 kit or 60 samples per S2 kit. Libraries were sequenced using 2×150 cycles on a NovaSeq 6000 (Illumina).

#### Pre-processing: alignment, annotation, and detection of gene fusions

The RNA sequencing data were processed as per the GATK 4.0 best practices workflow for variant calling, using a wdl and cromwell based workflow (https://gatk.broadinstitute.org/hc/en-us/sections/360007226651-Best-Practices-Workflows). This included performing quality control with Fastqc (v0.11.5) to calculate the number of sequencing reads and the insert size (Andrews S., 2010. FastQC: a quality control tool for high throughput sequence data, http://www.bioinformatics.babraham.ac.uk/projects/fastqc). Picard (v2.20.1) for RNA metrics output and MarkDuplicates (“Picard Tools.” Broad Institute. http://broadinstitute.github.io/picard/). The raw sequencing reads were aligned using Star (v2.7.0f) to GRCh38 and gencode version 31. Gene fusion detection was performed using Star fusion (v1.6.0) (Haas et al., 2019). Finally, expression counts were determined at exon and gene level using Subread Counts (Liao et al., 2019).

#### Processing of transcriptome data

Raw count tables were loaded into *R* (v4.0.2) and processed with the packages from the tidyverse. Count matrices were transformed into log2-scale transcripts per million (log2 TPM) tables and genes annotated with the same gene symbol merged.

Comparison of transcriptional profiles across samples (kidney and rhabdomyosarcomas) was performed via correlation performed in *R* using the base package.

### Bulk sequencing data availability

The process to make the bulk sequencing data (i.e., whole-genome sequencing and bulk mRNA sequencing) openly available at the European Genome-Phenome Archive (EGA) has been initiated with data upload completed and final curation pending.

### Growth curve experiments

Growth behavior in 3D sphere cultures prior to drug screenings (see below) was tested via performing growth curve experiments. Tumoroid cells were plated at different densities (typically between 250 and 4.000 viable cells per well) in 384-well round bottom ultra-low attachment spheroid microplates (Corning, cat no. 3830). To facilitate 3D sphere formation, cells were grown in BM1* medium without BME and after dispensing them into the plates, plates were spun (1.500 rpm, 5 min, slow break) to concentrate cells in the center of the wells. Plates were incubated at standard conditions and cells were granted a recovery period of 48 hours. Thereafter, readouts were performed using CellTiter-Glo 3D Cell Viability Assay (CTG3D, Promega, cat no. G9683) according to the manufacturer’s protocol at three time points: immediately (T0), 48 h or 72 h (T48 or T72), and 120 h (T120). Measured luminescence (via FLUOstar Omega, BMG Labtech) was averaged per readout per cell density and the corresponding background signal from medium was subtracted. Obtained measurements from T48/T72 and T120 were normalized to T0 to calculate the relative growth over the period of 120 h. For the following drug screening experiments (see below), a cell density was chosen that showed logarithmic growth behavior in growth curve experiments. Defined starting number of cells facilitated the establishment of a sphere and were optimized for intrinsic growth factor levels and space depletion in the well over the course of the experiment. Growth curve experiments were performed once per model but with at least ten technical replicates per number of cells plated.

### Drug screenings

For drug screenings, tumoroid models were processed according to the same protocol as for the growth curve experiments, with a number of cells seeded that was determined in those experiments (see above). Drugs, dissolved in DMSA or water, were added 48 hours after seeding of the tumoroid spheres fully automated via a robotics system: Up until 2019, this was facilitated via a Caliper Sciclone-Robotic Liquid Handler using a dilution of the drugs in medium and transferring this dilution to the cells by pipetting. From 2020 onwards, screenings were performed at the high-throughput screening (HTS) facility of the Princess Máxima Center with a Biomek i7 liquid handler, using the acoustic liquid handler Echo550 for direct drug transfers. Before the screening, the 384-well working plates containing the dissolved drugs are shaken (30 min, RT) and centrifuged (1 min, 1500 rpm). Tumoroid spheres were treated with a ten-fold dilution series of the drugs on the library plate (0.1 nM to 10 µM). Positive control samples were treated with DMSO, negative control samples with staurosporine (final concentration of 10 µM). Readouts were performed using CellTiter-Glo 3D Cell Viability Assay (CTG3D, Promega, cat no. G9683) according to the manufacturer’s protocol at T0 (before addition of drugs – control) and at T120 (120 h after addition of drugs – readout). Dose-response was estimated per drug and concentration in relation to the DMSO-treated cells (set to 100%) and empty controls (set to 0%). Quality of the screenings was approved after assessment of the cell growth (absorbance signal of T120 over T0), the negative, positive, and empty controls and, if applicable, the amount of variability between the duplicates.

Depending on the available number of cells at drug screening, a subset of models (RMS007, RMS109, RMS110, RMS000EEC, RMS000ETY, RMS000FLV, RMS000HQC, RMS000HWO, RMS000HWQ) could be screened in technical duplicates while for the other models (RMS006, RMS012, RMS013, RMS102, RMS108, RMS127, RMS335, RMS410, RMS444, RMS000CPU) only screening without technical duplicates was feasible. To test reproducibility, we analyzed the correlation between the technical duplicates in the tumoroid models for which those were available (Suppl. Fig. 5). As we observed a very high correlation there (*R* = 0.91), indicating a high reproducibility, we deemed it appropriate to combine the analysis of samples tested in duplicate and those not tested in duplicate given the scope of the assay was to assess whether we could obtain biologically meaningful results (any “hit” should have been further validated). RMS tumoroid models RMS000ETY and RMS000HWO were excluded from the analysis as they did not show an increase of CTG3D signal between T0 and T120 (Suppl. Table 2), indicating that these two models did not grow sufficiently under the screening conditions (while they displayed a minor increase in signal during the growth curve experiments).

Further data analysis was performed using *R* (v3.6.3). Area Under the Curve (AUC) values were calculated for every drug per tumoroid model and replicate (in the case of the tumoroid models with technical replicates) using the *auc* function of the MESS package (v0.5.6). In the case of RMS tumoroid models with technical duplicates, the two resulting AUC values per drug were correlated (using the *cor* function of base *R*) to calculate the correlation between replicates and plotted using the *plot* function of base *R* (see above). For the further downstream analysis, these replicate AUC values were averaged, resulting in a matrix with one AUC value per drug per RMS tumoroid model. On this matrix, unsupervised clustering was performed using the *get_dist* function from the factoextra package (v1.0.7) using the arguments “pearson” for RMS tumoroid models and “euclidean” for drugs to measure dissimilarity with further clustering using the *hclust* function from base *R* using the “average” argument for linkage. Data were visualized using the *heatmap.2* function from the gplots package (v3.0.3) using dendrograms generated from the established clustering. Principal component analysis (PCA) was performed using the *prcomp* function from base *R* with exclusion of the RMS tumoroid model RMS000FLV due to its outlier behavior. PCAs were visualized using the *fviz_pca_ind* and *fviz_pca_var* functions from the above-mentioned factoextra package.

### CRISPR/Cas9 knockout of *TP53* and functional evaluation

RMS012 tumoroid cells were kept under standard conditions (BM1* supplemented with 0.1 % BME) prior to the experiment. Upon reaching 70 % confluency, tumoroid cells were passaged as usual and seeded with high density into wells of a 24-well plate. After 24 h of recovery, transfection was performed: Nucleic acid–Lipofectamine 2000 complexes were prepared according to the standard Lipofectamine 2000 protocol (Invitrogen). Four µl of Lipofectamine 2000 reagent in 50 µl Opti-MEM medium (Gibco) and 1.5 µg of DNA (pSpCas9(BB)-2A-GFP control or sgRNA *TP53* plasmid in 50 µl Opti-MEM medium) were mixed, incubated for 5 min, and added to the cells. Plasmids were kindly shared by Jarno Drost (PMC, NL). For the plasmid sequence refer to Drost *et al*. 2015 (Drost et al., 2015). Twenty-four hours after transfection, transfection efficiency was evaluated using fluorescence microscopy, detecting GFP positive cells. Forty-eight hours after transfection, selection with nutlin-3 (10 µM) was started. Four days after the start of selection, first nutlin-3 resistant colonies could be detected in *TP53* knockout plasmid transfected cells whereas cells transfected with control plasmids died due to nutlin-3 exposure. Putative knockout cells were further expanded.

Knockout was confirmed using Western Blotting and Sanger sequencing of genomic DNA: For Western Blotting, snap frozen tumoroid cell pellets were lysed in Phosphatase-substituted RIPA buffer and run on a 10 % precast gel for P53 detection (BioRad). Protein levels of P53 (1:1.000, Santa Cruz DO-1 P53 antibody) were detected while Histone 3 (1:2.000, Abcam ab1791 Pan-H3) served as loading control. For visualization, secondary antibodies (goat anti-rabbit, BioRad 1706515 / goat anti-mouse BioRad 1706516) conjugated with horseradish peroxidase (HRP) were used together with ECL substrate (Perkin Elmer) on an imaginer (BioRad ChemiDoc). Western Blotting analysis was performed twice, and a representative blot is shown. For Sanger sequencing of genomic DNA, a snap frozen tumoroid cell pellet was lysed using DirectPCR Lysis Reagent for Cells (Viagen) according to the manufacturer’s protocol. PCR amplification was performed using Phusion High-Fidelity DNA Polymerase (New England BioLabs) using the following primers: forward 5’-CCCATCTACAGTCCCCCTTG-3’, reverse 5’-CAGGAAGCCAAAGGGTGAAGA-3’. PCR products were cleaned up and concentrated using a DNA Clean-up and Concentration Kit (Zymo Research) according to the manufacturer’s protocol. Gel electrophoresis indicated the presence of a specific product which was sent for Sanger sequencing using the following primers: forward 5’-TGGTTCACTGAAGACCCAGG-3’, reverse 5’-GAAGTCTCATGGAAGCCAGCC-3’. Obtained sequences were aligned and inspected using the Benchling browser tool (https://www.benchling.com/). Furthermore, sequencing data was submitted for TIDE (Tracking of Indels by Decomposition, http://tide.nki.nl) analysis to infer the composition of Indels in the knockout population (Brinkman et al., 2014).

For detection of DNA double strand breaks, induction of γH2AX was measured using Western Blotting. Fusion-negative embryonal rhabdomyosarcoma cell line RD (kindly shared by Jan Molenaar, PMC, NL) served as a control (either untreated or treated with 1 µM, 5 µM, or 10 µM of staurosporine (Sigma) for 24 h). RD cells were cultured under conventional conditions as detailed by the American Type Culture Collection (ATCC) with regular testing for mycoplasma contamination. For Western Blotting, snap frozen tumoroid (RMS012 *TP53* KO) or tumor (RD) cell pellets were lysed in Phosphatase-substituted RIPA buffer and run on a 15% self-cast gel for γH2AX (p.S139) detection. Protein levels of γH2AX (p.S139, 1:2.000, Abcam ab26350 antibody) were detected while GAPDH (1:1.000, Abcam ab9485 antibody) served as loading control. Visualization was performed as described above for the confirmation of the knockout. As above, Western Blotting analysis was performed twice, and a representative blot is shown.

To assess differential response to prexasertib (MedchemExpress), RMS012 tumoroid models (*TP53* wildtype and *TP53* knockout) were processed according to the same protocol as for the growth curve experiments (see above), with 500 cells seeded per well. Prexasertib (dilution series from 200 nM to 0.78125 nM) was added 48 hours after seeding of the tumoroid spheres. Control samples were treated with DMSO. Readouts were performed using CellTiter-Glo 3D Cell Viability Assay (CTG3D, Promega, cat no. G9683) according to the manufacturer’s protocol at T72 (72 h after addition of drugs). Dose-response was estimated per model and concentration in relation to the DMSO-treated cells (set to 100%). The experiment was performed three times. Further data analysis was performed using *R* (v3.6.3). A sigmoidal fit for the dose-response curve was calculated per replicate for both the knockout and wildtype sample with a three-parameter log-logistics function using the drc package (v3.0-1) (Ritz et al., 2015). The statistical significance of the differences in fitted IC_50_ values between knockout and wildtype were obtained using a two-sided t-test. For visualization purposes a three-parameter sigmoidal fit per model (not per replicate) was used.

### Image post-processing and figure preparation

Microscopy images from RMS tumoroid IHC and H&E stainings as well as images from Western Blotting were post-processed according to good scientific practice with Adobe Photoshop 2021 and Fiji (v2.0.0-rc-69/1.52i) (Schindelin et al., 2012). Images from original RMS tumors (H&E and IHC) were not processed. Figures were prepared using Adobe Illustrator 2021.

### Code availability

Code is made openly available on https://github.com/teresouza/rms2018-009.

## Supplemental items legends

**Suppl. Fig. 1:**
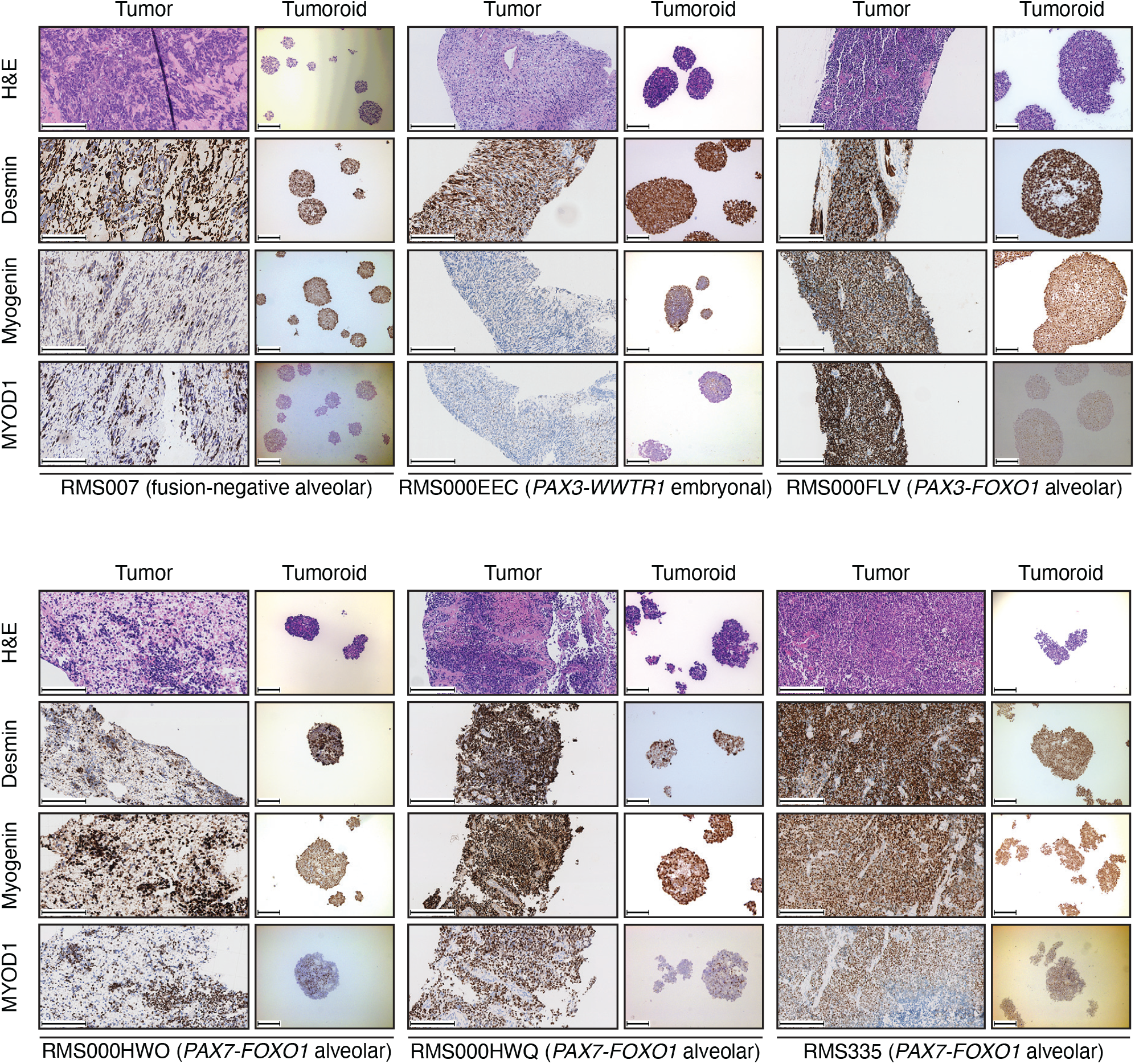
Morphological (via H&E) and immunohistochemical (IHC) comparison of RMS tumors and derived RMS tumoroid models shows retained marker protein (Desmin, Myogenin, and MYOD1) expression and cellular morphology. Scale bars equal 200 µm.

**Suppl. Fig. 2:**
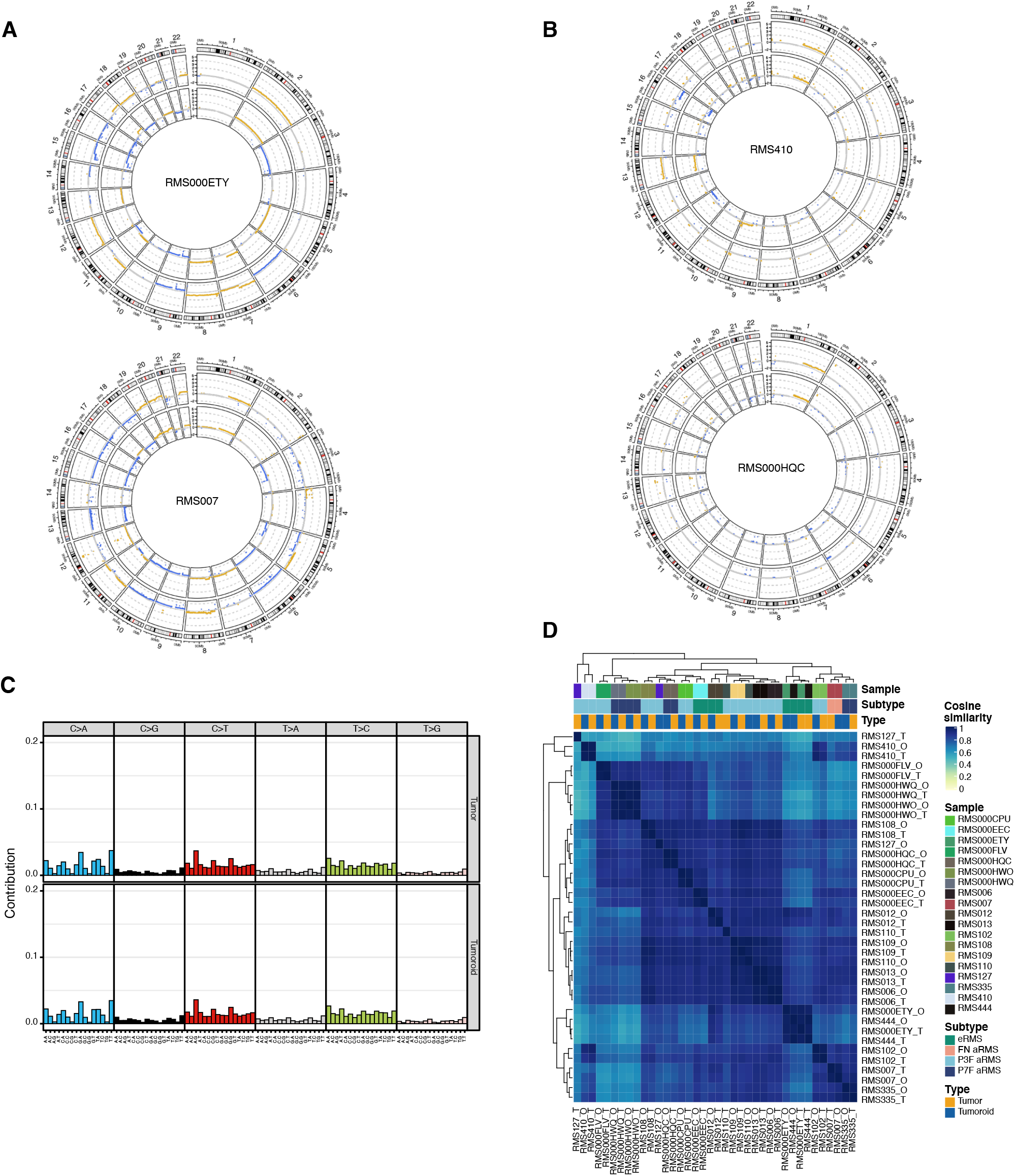
(A) Examples of circular copy number plots of fusion-negative RMS with embryonal (RMS000ETY) or alveolar (RMS007) histology. Outer circle depicts the tumoroid model, inner circle depicts the tumor. (B) Examples of circular copy number plots of fusion-positive RMS with alveolar histology and *PAX3-FOXO1* fusion (RMS410) or *PAX7-FOXO1* fusion (RMS000HQC). Outer circle depicts the tumoroid model, inner circle depicts the tumor. (C) Contribution of averaged single base substitution (SBS) profiles for RMS tumors (upper row) and tumoroid models (lower row). (D) Clustered correlogram of SBS profiles detected in RMS tumors (T) and tumoroid models (O).

**Suppl. Fig. 3:**
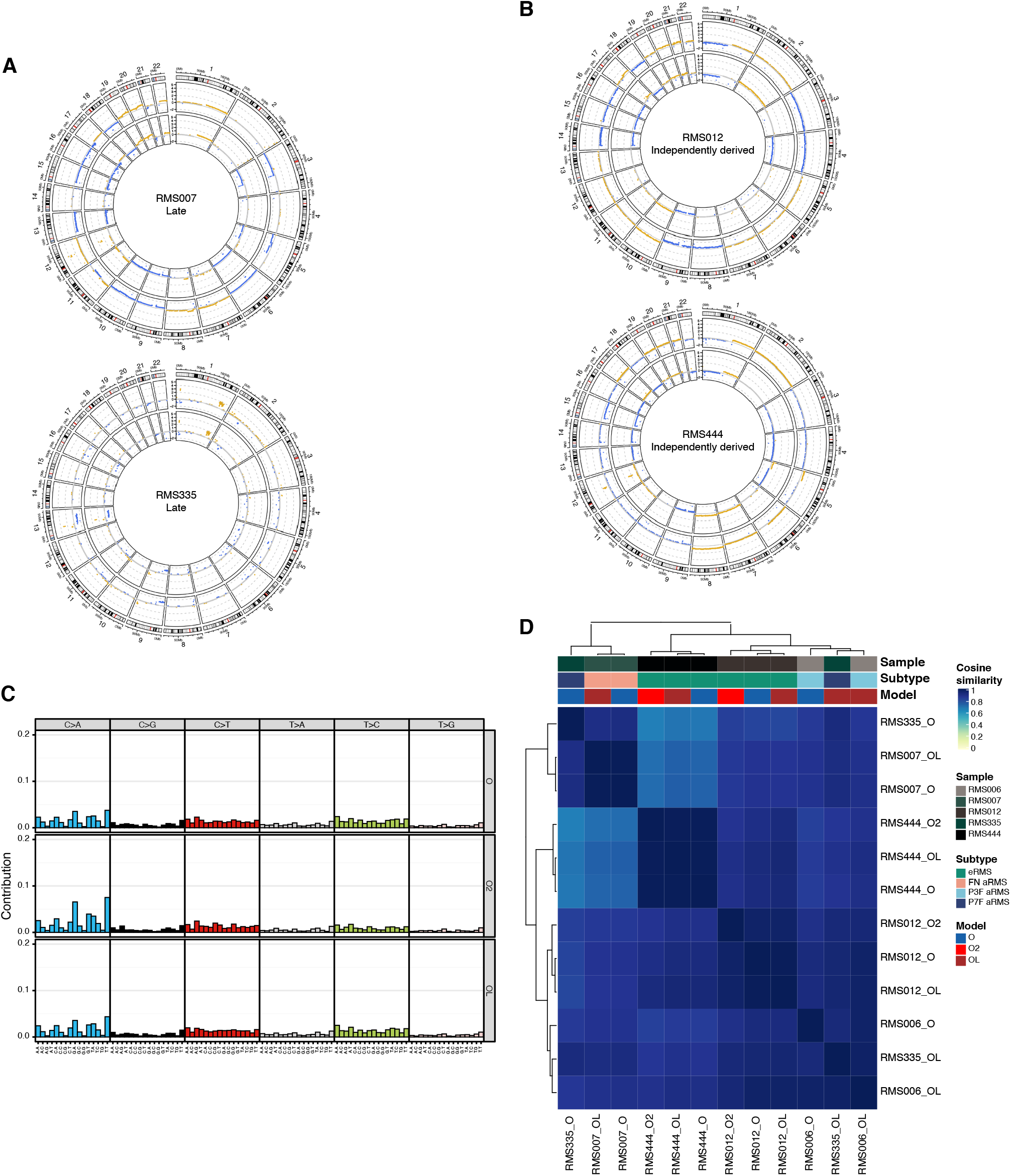
(A) Circular copy number plots of standard and late passage RMS tumoroid models of a fusion-negative alveolar (RMS007) and a *PAX7-FOXO1* positive alveolar (RMS335) RMS. Outer circle depicts the standard passage tumoroid, inner circle depicts the late passage tumoroid model. (B) Circular copy number plots of standard passage and independently derived RMS tumoroid models of two fusion-negative embryonal RMS (RMS012 and RMS444). Outer circle depicts the standard passage tumoroid, inner circle depicts the independently derived tumoroid model. (C) Contribution of averaged single base substitution (SBS) profiles for standard (O), independently derived (O2), and late passage (OL) RMS tumoroid models. (D) Clustered correlogram of SBS profiles detected in standard (O), independently derived (O2), and late passage (OL) RMS tumoroid models.

**Suppl. Fig. 4:**
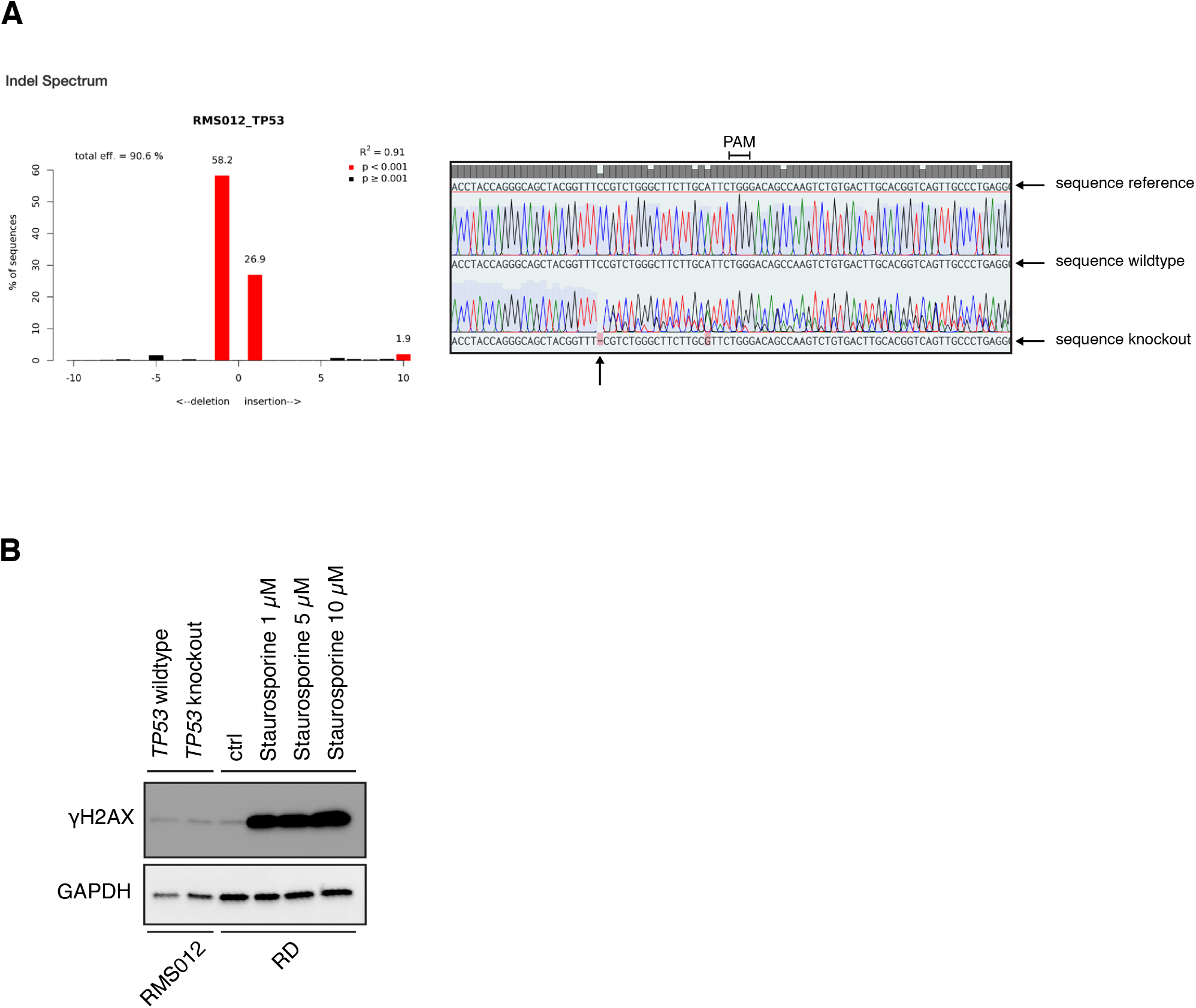
(A) Left panel: Inferred Indel Spectrum from Sanger sequencing using the TIDE tool (https://tide.nki.nl). Right panel: Sanger sequencing around Cas9 cut site (arrow at the bottom) in *TP53* gene. Protospacer Adjacent Motif (PAM) sequence indicated. (B) Western Blotting analysis of γH2AX (Ser-139) as proxy for DNA double-strand breaks in untreated RMS012 tumoroid cells (*TP53* wildtype and *TP53* knockout). RMS cell line RD treated with staurosporine served as positive control while GAPDH served as loading control.

**Suppl. Fig. 5:**
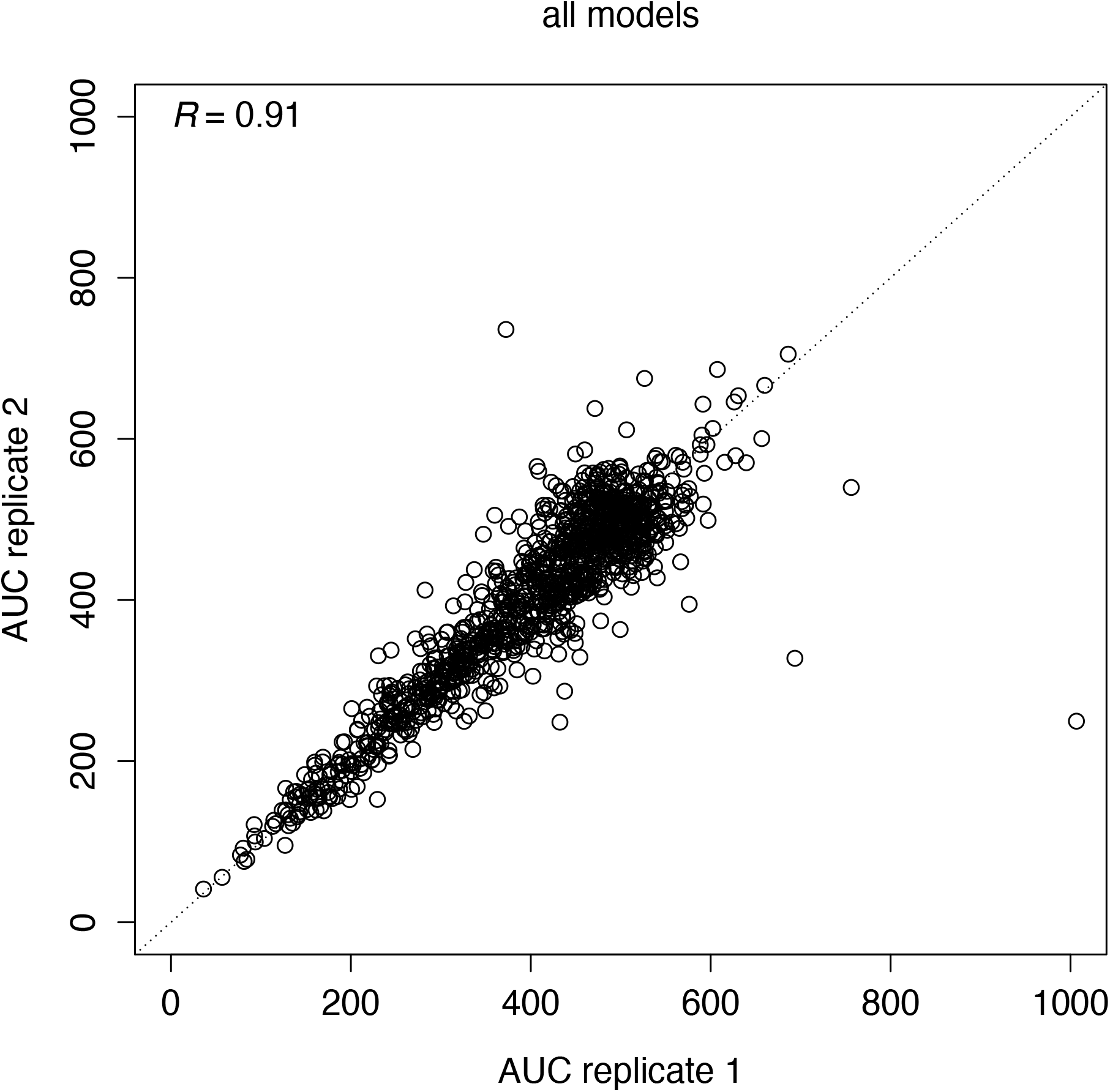
Correlation analysis of Area Under the Curve (AUC) values from technical replicates of drug screenings of RMS007, RMS109, RMS110, RMS000EEC, RMS000FLV, RMS000HQC, and RMS000HWQ. Overall, the correlation coefficient (*R*) is 0.91.

## Table legends

**Suppl. Table 1:** Clinical annotation RMS tumoroid model cohort.

**Suppl. Table 2:** Relative growth of vehicle treated control tumoroid cells during drug screenings.

## Notes

https://rmsdatabiobank.shinyapps.io/app1_3/

### Competing Interest Statement

The authors have declared no competing interest.

